# Queuosine modification mediates cold-active growth in *Shewanella glacialimarina*

**DOI:** 10.64898/2026.05.06.723185

**Authors:** M. Suleman Qasim, Jenni K. Pedor, Alisa Willman, Johannes Merilahti, Otto Kauko, Nina Sipari, L. Peter Sarin

## Abstract

Efficient protein synthesis in cold-active bacteria requires precise coordination within the translation machinery to overcome the kinetic challenges imposed by growth at near-freezing temperatures (< 5 °C). Post-transcriptional modifications (PTMs) on transfer RNA (tRNA)—particularly those located at the wobble position 34—are central to this coordination as they regulate decoding speed and fidelity. Here, we show that queuosine (Q) modification in the cold-active marine bacterium *Shewanella glacialimarina* TZS-4_T_ is dynamically regulated in response to bacterial growth and environmental conditions. Importantly, we demonstrate that Q levels are modulated in a tRNA isoacceptor-specific manner during cold-active growth, while Q-deficiency produces a cold-sensitive phenotype that underscores the functional importance of Q modification. Proteomic analysis of the Q-deficient Δ*tgt* mutant revealed that tRNA^His^ Q-hypomodification activates the histidine biosynthesis pathway, whereas the concomitant phosphate-starvation-like response reflects a general consequence of global disruption of Q-modified tRNAs. Consequently, we propose a model where Q modification maintains efficient codon decoding and protein quality control at near-freezing temperatures, whereas loss of Q destabilizes codon decoding and ultimately compromises protein homeostasis.

## INTRODUCTION

It is estimated that ∼80 % of Earth’s biosphere is subjected to temperatures that are permanently or periodically below 5 °C, which necessitates specialized adaptation strategies for life to survive (Russell, 1990; De Maayer *et al*., 2014). In cold habitats (< 10 °C), psychrophilic organisms are challenged with reduced membrane fluidity, an increase in water viscosity, and low kinetic rates of metabolic reactions. Psychrophiles and psychrotrophs mitigate these effects by enriching their cell membrane with unsaturated fatty acids (Allen *et al*., 1999; Nichols *et al*., 1993), weakening intramolecular interactions within enzymes (D’Amico *et al*., 2006; Johns & Somero, 2004) and inducing cold-shock proteins to promote mRNA translation (Berger *et al*., 1996). While these adaptations in membrane composition, enzyme dynamics, and gene regulation are well established, our understanding of how the translation machinery—a vital energy-intensive process—adapts to cold is largely based on a few model organisms and remains poorly characterized.

At the centre of translation lies transfer RNA (tRNA)—it serves both as an adapter that decodes mRNA codons and as a carrier that delivers the cognate amino acid to the nascent polypeptide chain. The functional diversity of tRNA, its intricate interactions with multiple components of the translation machinery, and its modulation by over 150 post-transcriptional modifications (PTMs) make it a compelling target to study in cold-adapted microorganisms. PTMs include multiple chemistries, such as methylation, acetylation, deamination, hydroxylation, selenylation, isomerization, and amino acid or sugar conjugation (Sordyl et al., 2026). In the case of thermophilic organisms, specialized PTMs (e.g. 5-methyl-2-thiouridine, m^5^s^2^U) are crucial for modulating the physiochemical properties of tRNA and ensuring optimal function (Watanabe *et al*., 1976, 1974). These modifications govern critical aspects of tRNA biology, including folding, thermal stability, and decoding efficiency (reviewed in Ohira & Suzuki, 2024). Their functional roles are primarily dependent on their location and the chemical properties of the modifications. PTMs on the tRNA body (D loop, TΨC loop, and the variable loop) primarily contribute to its stability (Motorin & Helm, 2010), whereas modifications within the anticodon stem loop (ASL) regulate decoding by either enhancing or limiting the recognition of specific codons (reviewed in Suzuki, 2021) or by preventing frameshifting (Urbonavičius *et al*., 2001). Such epitranscriptomic regulation becomes especially important during stress conditions, offering a dynamic regulatory framework to overcome significant constraints that repress translation and cellular activity. For example, in psychrophilic microorganisms, dihydrouridine (D) in the D loop promotes the C2’-endo conformation of tRNA and disrupts base-stacking, thereby increasing local flexibility of tRNA. In addition, loss of *trmE*, the enzyme responsible for generating 5-methyl-aminomethyluridine (mnm^5^U) at wobble position 34, results in a cold-sensitive phenotype due to transient blocks in translation caused by frameshifting (Singh *et al*., 2009).

To investigate the dynamic reprogramming of tRNA modifications under cold stress, we utilized *Shewanella glacialimarina* TZS-4_T_, a cold-active marine bacterium originally isolated from Baltic Sea ice. We have previously shown that *S. glacialimarina* is uniquely adapted to low temperatures—it grows between 0 °C and 25 °C with an optimum at 15 °C, and it enhances membrane fluidity at low temperatures by increasing the abundance of polyunsaturated fatty acids (Qasim *et al*., 2021). In a subsequent study, we further demonstrated that queuosine (Q) modification increases during viral infection, which coincides with the expression of a viral transcript with a significant UAC codon preference (Lampi *et al*., 2023). However, the effect of physiological stressors on the tRNA modification landscape of *S. glacialimarina*, as well as their subsequent impact on translation and thereby survival of the bacteria remains unexplored. It has previously been shown that tRNA modifications can dynamically regulate the translation of codon-biased genes, which may be essential for adaptation during stress conditions (Endres *et al*., 2015). Yet, it is unclear whether such codon-biased translation contributes to growth in cold-active bacteria, as our current understanding of prokaryotic tRNA modifications largely stems from a few mesophilic model organisms, such as *Escherichia coli* and *Mycoplasma capricolum*. While recent work in *Bacillus subtilis* and *Vibrio cholerae* has broadened our knowledge to other bacterial genera (de Crécy-Lagard *et al*., 2020; Fruchard *et al*., 2025), studies on tRNA PTMs and their effect on translation at low temperatures remain scarce.

Here, we employed an integrated multi-omics approach to comprehensively characterize how the tRNA modification landscape in *S. glacialimarina* is regulated under cold stress. We demonstrate that cold temperatures (0 °C and 5 °C) trigger an increase in Q modification specifically on tRNA^Asp^, whereas Q levels on tRNA^His^ and tRNA^Tyr^ are reduced. Importantly, loss of Q (Δ*tgt* mutant) results in a pronounced cold-sensitive phenotype driven by two distinct stress factors: (i) reduced uptake of inorganic phosphate (P_i_), possibly due to inefficient translation of the PstS outer membrane phosphate transporter protein, and (ii) impaired tRNA^His^ function. In this scenario, translational inefficiency leads to a phosphate-starvation-like phenotype, which in turn activates expression of the Pho regulon and associated phosphatase enzymes. Likewise, compromised tRNA^His^ functionality, either by mischarging or decoding defects, induces the histidine biosynthesis pathway. Hence, the synergistic effect of these stress responses results in an increased metabolic burden and a heightened sensitivity to cold temperatures. Consequently, we propose that Q modification acts as a protein quality control factor—when present, it upholds efficient decoding at low temperatures but when lost, codon-anticodon interactions become destabilized and protein homeostasis is perturbed.

## RESULTS

### Combined tRNA-seq and UPLC/MS uncovered 17 distinct tRNA modifications in *S. glacialimarina*

*S. glacialimarina* encodes 92 tRNA genes grouped into 33 tRNA isoacceptor families, carrying distinct anticodons for the same amino acid **(Supplementary Table T1)**. Full-length tRNA sequencing using MarathonRT revealed sequence heterogeneity within several of these isoacceptor families, uncovering 43 uniquely expressed tRNA species that encompass 33 isoacceptor families **(Supplementary Table T2)**. Based on the genome annotation, we identified 37 putative tRNA modification enzymes predicted to introduce 24 distinct modification types across the tRNA pool **(Supplementary Table T3)**, including for example various methylations of both the nucleobase and sugar moiety, thiolation, pseudouridinylation (Ψ), D, and Q. To experimentally determine the predicted tRNA modification profile, we performed broad-range UPLC/MS analysis (Gregorova *et al*., 2021) of *S. glacialimarina* bulk tRNA, which allowed us to detect and identify 17 of the 24 predicted modifications **(Figure 1)**. The identity of 2’-O-methylcytidine (Cm), N6-methyladenosine (m^6^A), 5-methyluridine (m^5^U), inosine, 1-methylguanosine (m^1^G), 7-methylguanosine (m^7^G), and 2’-O- methylguanosine (Gm) was established based on available ribonucleoside standards. However, D, 2-lysidine (k^2^C), 5-methylaminomethyl-2-selenouridine (mnm^5^se^2^U) and N6-isopentenyladenosine (i^6^A) could not be detected **(Supplementary Data File 1)**, most likely due to their low abundance and/or weak ionization intensity, which caused the signal to fall below the detection limit. The remaining 11 PTMs were inferred by ionization pattern matching to previously reported data in Modomics (Sordyl *et al*., 2026). To further complement this dataset, we performed tRNA sequencing and examined misincorporation signatures generated using MarathonRT reverse transcriptase in the mim-tRNAseq pipeline (Behrens *et al*., 2022; Behrens & Nedialkova, 2022), which provided positional information of specific modifications on individual tRNA isoacceptors. Hence, the remaining 11 PTMs were additionally verified based on the presence or absence of (a) corresponding tRNA modification enzyme gene(s), supporting data from mim-tRNAseq, and/or previously reported ionization patterns **(Figure 1)**.

**Figure 1.**
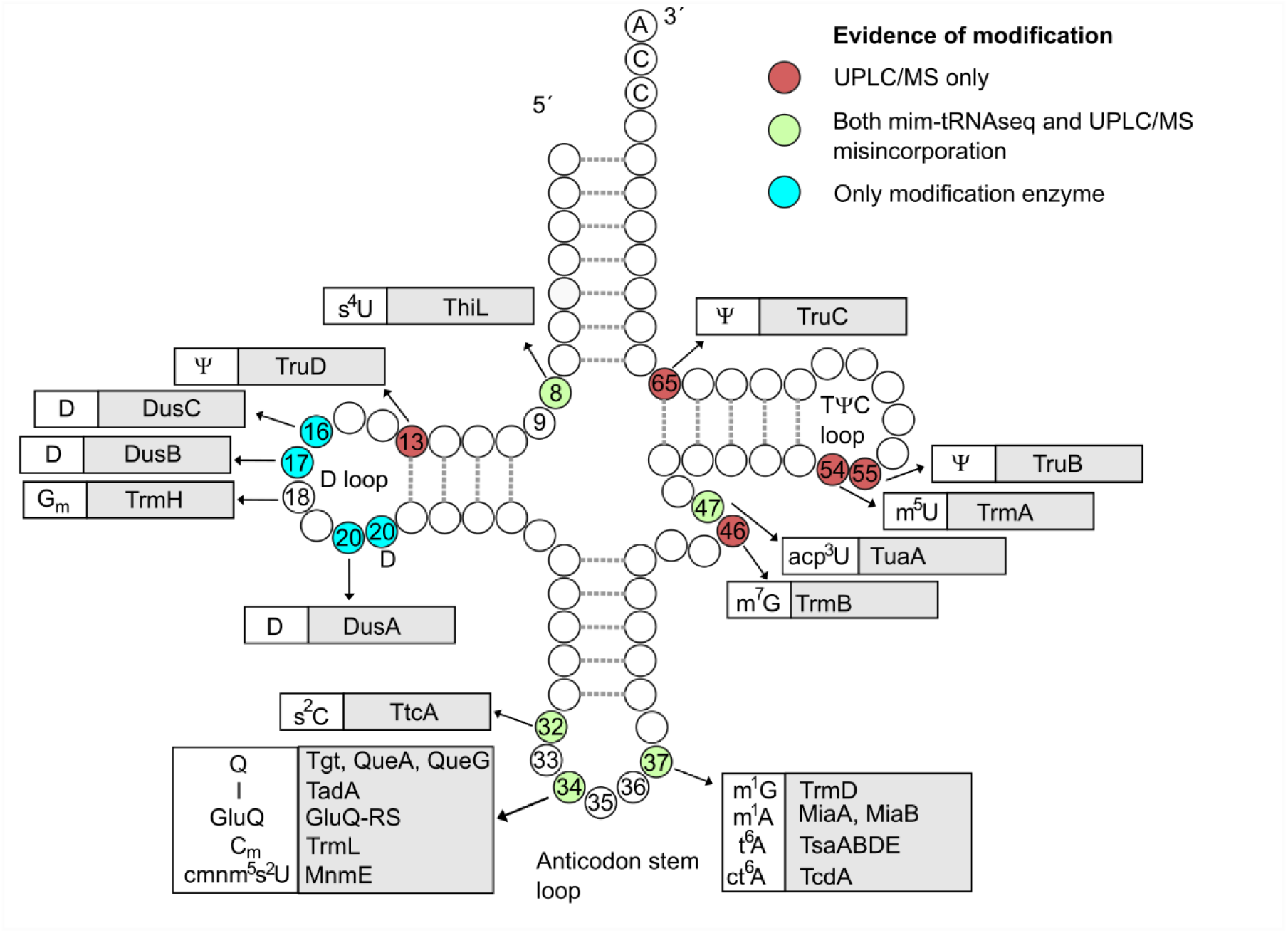
Post-transcriptional tRNA modifications in *S. glacialimarina* TZS-4_T_. Schematic representation of a mature transfer RNA showing modification sites and identities. The annotations are based on (i) UPLC-MS data, (ii) combined mim-tRNAseq misincorporation and UPLC-MS data, and (iii) the presence of modification enzyme in the genome. The colour scheme shown at the top right depicts the source of evidence for each modification site. Enzymes responsible for catalysing each tRNA modification are shown adjacent to their respective modification site(s) (box with grey background).

Moreover, the mim-tRNAseq results also revealed major misincorporation events at positions 8, 34, and 47 in multiple tRNAs, which have previously been described as 4-thiouridine (s^4^U), inosine, and 3-(3-amino-3-carboxypropyl)uridine (acp^3^U) modification sites, respectively **(Supplementary Table T4)**. These PTMs disrupt Watson–Crick base pairing and generate a characteristic misincorporation signature during reverse transcription, which is consistent with our observations at the aforementioned positions in full-length tRNA transcripts. In the mim-tRNAseq dataset, the most widespread misincorporation was observed at position 8 for the predicted s^4^U modification, which appears in 17 out of 43 isoacceptors. Similarly, misincorporation at position 47 (predicted acp^3^U) was noted for 9 isoacceptors. We also identified additional misincorporation sites at positions 20, 22, 34, 37, and 46, which most likely arise from the presence of methyl groups, namely Gm, Cm, m^1^G, and m^7^G **(Supplementary Table T4)**. However, as the proposed positions for these PTMs rely on information derived from other model organisms and the mim-tRNAseq misincorporation patterns, the exact location of these PTMs cannot be determined without additional verification, such as LC-MS-based oligonucleoside sequencing.

### Post-transcriptional modification is linked to the bacterial growth phase

To investigate the temporal dynamics of *S. glacialimarina* tRNA modifications at standard liquid culture conditions (25 % rMB, 15 °C), we performed a time course study and collected cells for bulk tRNA isolation at hourly intervals between 3 h and 8 h **(Figure 2A)**. The extracted tRNA pools were analysed by UPLC/MS and the fold change in PTM abundance (relative to the 3 h timepoint) was determined across the 4_–_8 h timepoints **(Figure 2B)**. Next, we analysed the tRNA modification levels to elucidate temporal changes, which uncovered four main trends in PTM dynamics to which the modifications were grouped: (1) initial increase followed by reduction, e.g. Q **(Figure 2C)**; (2) sustained accumulation, e.g. Cm **(Figure 2D)**; (3) transient fluctuations, e.g. acp^3^U (**Figure 2E)**; and (4) consistent decrease, e.g. inosine (I) (**Figure 2F)**.

**Figure 2.**
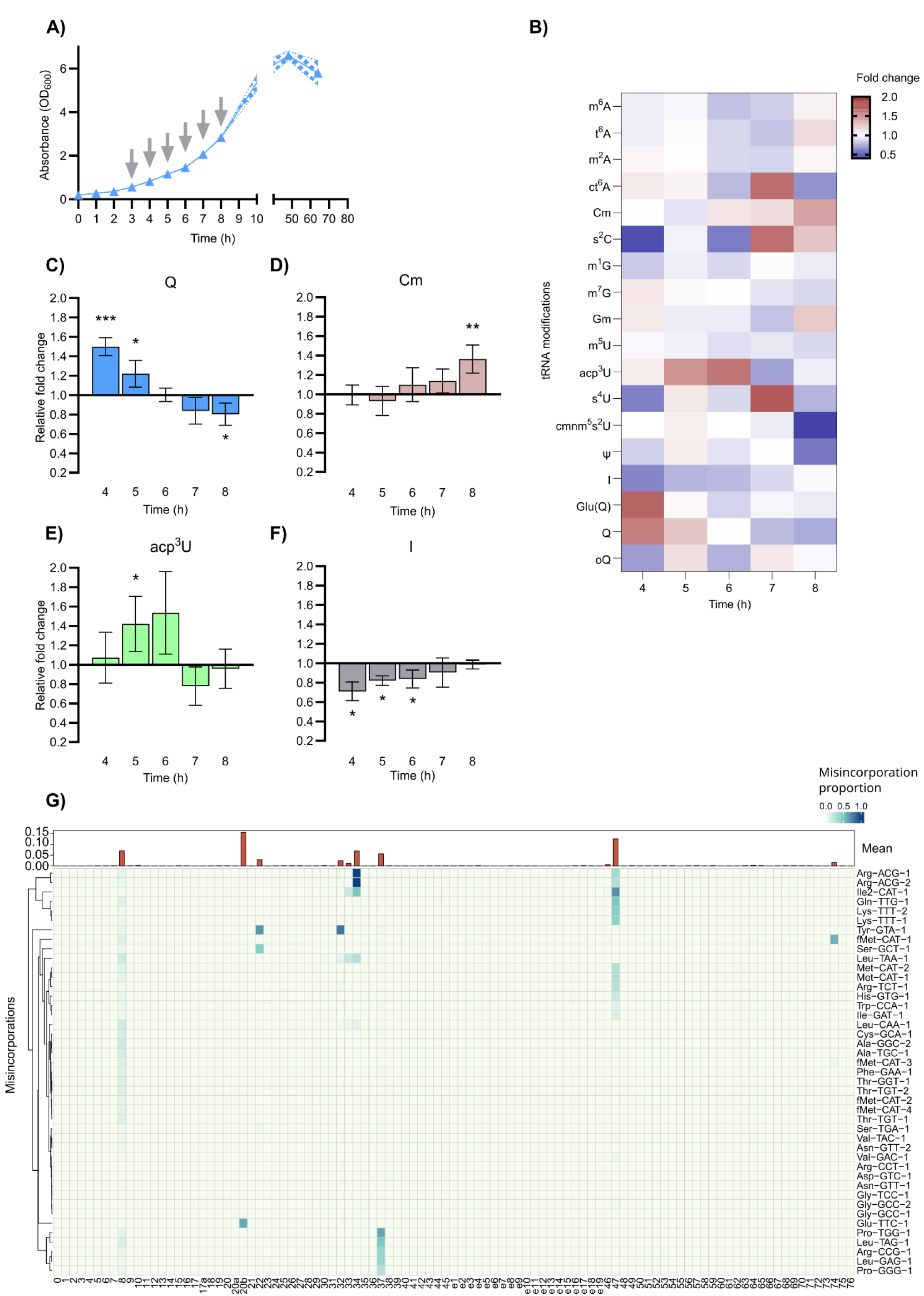
Dynamics of the *S. glacialimarina* tRNA modification landscape during exponential growth. A) Representative growth curve assay at 15 °C (n = 4). Grey arrows indicate sample collection timepoints. B) UPLC/MS analysis of tRNA modification levels (fold change relative to PTM intensity at 3 h post-inoculation) at selected exponential growth phase timepoints (n = 3). C–F) Bar plots of selected PTMs grouped according to modification dynamics (fold change trend): C) decreasing, D) increasing, E) transient fluctuations, or F) initial decrease and recovery. Error bars depict standard deviation (n = 3) and *p*-values were calculated using one-sample Student’s *t-*test with significance represented as * *p* < 0.05, ** *p* < 0.005, *** *p* < 0.0005. G) Overview of tRNA isoacceptor-specific misincorporation patterns following mim-tRNA-seq analysis using MarathonRT. The concatenated mean misincorporation levels are shown at the top of the panel (n= 4).

For group 1, we observed that Q levels peaked at 4 h (1.5-fold increase) and remained elevated at 5 h, which corresponds to the early exponential growth phase **(Figure 2C)**, whereas a slight decrease (0.8-fold) was observed at 7–8 h **(Figure 2C)**. This shift might indicate an adaptive modulation of wobble anticodon modification in response to changing physiological conditions. Interestingly, at standard growth conditions (15 °C) we also detected epoxyqueuosine (oQ)—the final precursor to Q—which displayed no discernible pattern (fold change 0.7_–_1.1) **(Supplementary Data File 1)**. Since oQ is efficiently converted to Q by the epoxyqueuosine reductase (QueG) (Miles et al, 2011), a growth-related increase in oQ levels may reflect reduced QueG activity and impairment of Q synthesis at its final step. Additionally, we also detected parent and product ions identical to glutamyl-queuosine Glu(Q) (observed *m/z* = 539.17 and *m/z* = 407.10, predicted *m/z* = 539.21 and *m/z* = 407.10) at RT 2.14 min for all the samples. This sugar conjugated Q is a stable derivative and widely known to be distributed across multiple bacterial genera on position 34 of tRNA^Asp^ (Blaise *et al*., 2004; Salazar *et al*., 2004). In our analysis **(Figure 2B),** Glu(Q) exhibited a pronounced 1.6-fold increase at 4 h, before returning to baseline level from 5 h to 8 h. This increase in Glu(Q) corresponds with the increase in Q at 4 h timepoint, suggesting a generalized phenomenon during early growth. Furthermore, we also identified N6-threonylcarbamoyladenosine (t^6^A; *m/z* = 413.14) and cyclic N6-threonylcarbamoyladenosine (ct^6^A; *m/z* = 395.13) based on their unique ionization profiles. Unlike oQ, both t^6^A and ct^6^A are stable chemical modifications. The conversion to cyclic form is accomplished by tRNA threonylcarbamoyladenosine dehydratase A (found in *S. glacialimarina*) by an ATP-dependent dehydration process (Miyauchi *et al*., 2012). While t^6^A abundance remained relatively stable across all timepoints, ct^6^A increased by 1.5-fold at the 7 h timepoint **(Supplementary Data File 1)**.

On the other hand, Cm and acp^3^U exhibited a subtle increasing trend **(Figure 2D, E).** Cm reached its highest level at 8 h (1.4-fold increase), whereas acp^3^U displayed an upward trend with high variability and a significant increase at the 5 h timepoint. Cm is known to occur at the wobble position of tRNA^Leu^ (CAA), whereas acp^3^U is found at position 47 of tRNA^lle^, tRNA^Arg^, tRNA^Lys^, tRNA^Met^, and tRNA^Trp^. Indeed, we observed distinct misincorporation pattern for these tRNAs at the respective sites, suggesting the presence of nucleoside modification **(Figure 2G)**. Lastly, inosine (I) modification presented a decreasing trend from 3 h to 6 h followed by a modest recovery to baseline levels at 7 h and 8 h **(Figure 2F)**. This modification site was also detected in the tRNA-seq dataset for tRNA^Arg^ (ACG), appearing as a misincorporation event at position A34 **(Figure 2G)**. Unlike adenosine, which pairs only with CGU, the structural flexibility of inosine allows pairing with C-, U-, and A- ending codons, thereby facilitating efficient decoding of CGU, CGC, and CGA (Crick, 1966). Despite this, *S. glacialimarina* showed a strong bias in the coding sequences (CDS) towards the CGU codon (RSCU = 2.44), which represents 40 % of all arginine codons. This suggests that although inosine is functionally important, its higher abundance may not be essential at the early growth phase.

### Oxidative and osmotic stress elicit divergent changes to the tRNA modification landscape

Next, we set out to investigate how *S. glacialimarina* tRNA modifications respond to environmental pressures—namely oxidative and osmotic stress—that marine bacteria encounter in their natural habitats. First, we induced a transient oxidative stress by applying 0.5 mM of hydrogen peroxide (H_2_O_2_) at OD_600_ = 0.5 (Qasim *et al*., 2021) and analysed PTM levels after 2 h, 4 h, and 6 h of H_2_O_2_ exposure. The initial responses were pronounced, and we observed a general decrease in most PTMs at 2 h, as highlighted by Q **(Figure 3 A)**, 2-methyladenosine (m^2^A), m^6^A, Gm, and m^7^G **(Figure 3 B)**, as well as t^6^A and ct^6^A **(Supplementary Data File 1)**. A similar response was observed for thiolation; 2-thiocytidine (s^2^C) decreased but s^4^U could not be detected, presumably since the signal intensity at 2 h was below the detection limit **(Figure 3C)**. Nevertheless, the effect of H_2_O_2_ exposure was largely transient, as most modifications returned to near baseline levels after 4 h and 6 h from exposure **(Figure 3 A**_–_**C)**. Then, we performed growth assays in high salinity conditions and noted a reduced growth rate in the presence of 55.5 g/L NaCl, which also constitutes the maximum NaCl tolerance for *S. glacialimarina* **(Figure 3 D)**. Interestingly, osmotic stress elicits a markedly different response compared to oxidative stress and is characterized by increased PTM levels. This is particularly evident for PTMs located at the main body of the tRNA, such as acp^3^U, Ψ, s^2^C and s^4^U **(Figure 3E, F)**. Indeed, we observed that acp^3^U and Ψ levels were more than 2-fold higher at high salinity conditions **(Figure 3E)**, while s^2^C and s^4^U levels increased by ≈1.5-fold **(Figure 3F)**. Since acp^3^U and Ψ are primarily located on the TΨC loop, this suggests that they stabilize the tRNA structure in high salinity conditions (Davis, 1995; Takakura *et al*., 2019). This may be further aided by s⁴U at position 8, which is predicted to engage with the D loop and act cooperatively with the TΨC loop to preserve the canonical L-shaped architecture of tRNA (Neumann *et al*., 2014; Kimura & Waldor, 2019). Finally, our UPLC/MS analysis also revealed a > 2-fold increase in oQ and Q levels, respectively, along with a modest rise in Glu(Q) (≈1.5-fold) **(Figure 3G)**. Hence, as oQ is a precursor of Q, this observed increase suggests a possible impairment in the Q biosynthesis pathway during osmotic stress.

**Figure 3.**
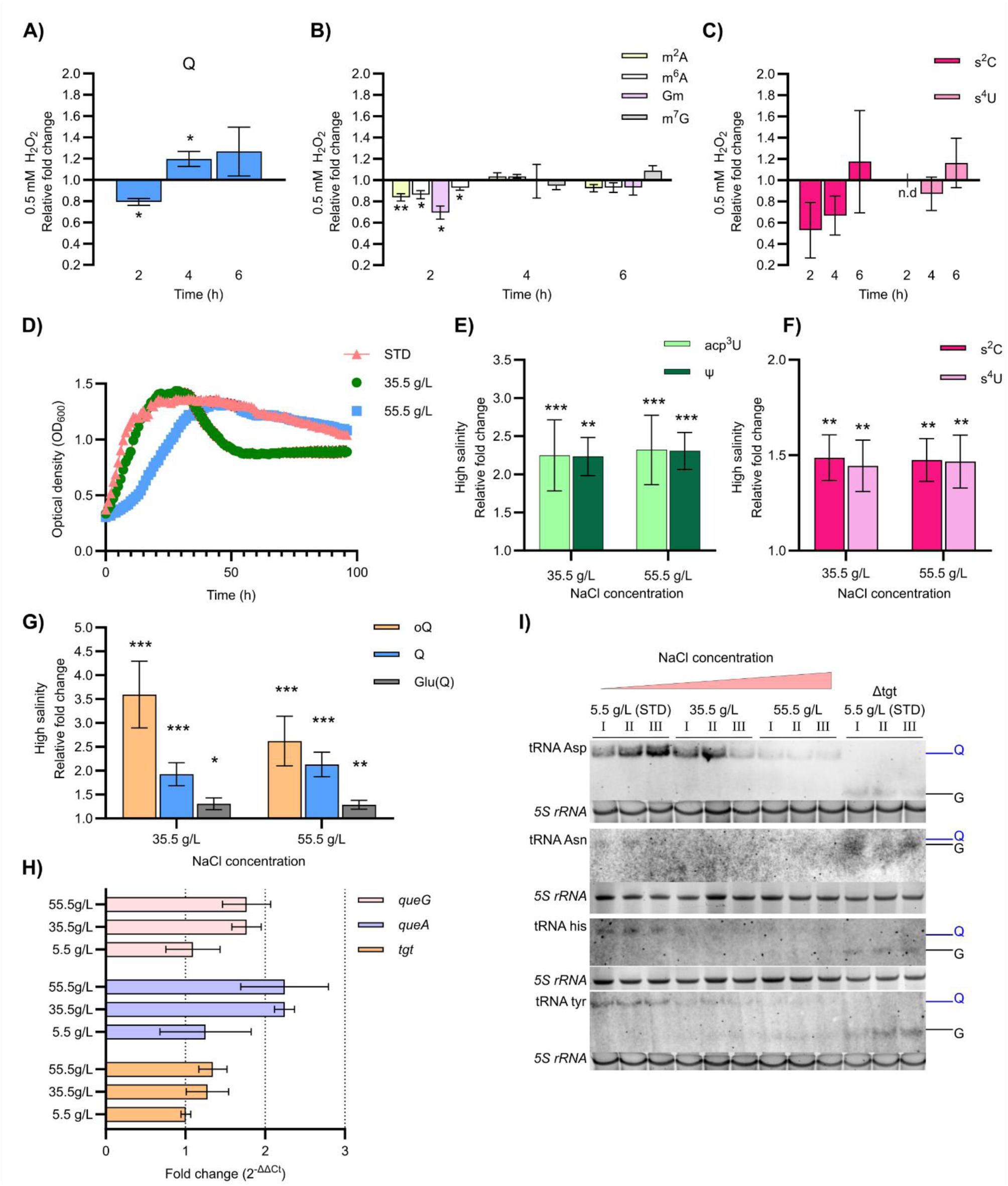
Stress-induced tRNA modification changes in *S. glacialimarina*. A-C) Bar plot representation of relative fold change in PTM levels during oxidative stress for selected modifications at 2 h, 4 h, and 6 h post-exposure to 0.5 mM H_2_O_2_. Relative fold change determined as the normalized PTM intensity for stressed vs. non-stressed (no H_2_O_2_) samples at each timepoint. Error bars denote the standard deviation and *p*-values, calculated within the sample size n = 4 using one-sample Student’s *t-*test with significance represented as * *p* < 0.05, ** *p* < 0.005, *** *p* < 0.0005. D) Growth of *S. glacialimarina* at high salinity (35.5 g/L and 55.5 g/L NaCl) and STD (5.5 g/L NaCl) growth conditions. The shaded area denotes the standard deviation within the curve. E-G) Bar plots for selected PTMs at high salinity (35.5 g/L and 55.5 g/L NaCl), depicted as fold change relative to STD (5.5 g/L NaCl) at the early exponential growth phase (OD_600_ = 0.8). Error bars denote the standard deviation and *p*-values, calculated within the sample size n = 4 using one-sample Student’s *t-*test with significance represented as * *p* < 0.05, ** *p* < 0.005, *** *p* < 0.0005. H) Gene expression analysis of transcripts for Q tRNA modification pathway members at STD (5.5 g/L NaCl) and high salinity (35.5 g/L and 55.5 g/L NaCl) conditions. The expression analysis was performed using the ΔΔ_CT_ method with the *gyrA* gene as reference. Error bars denote the standard error (n = 3). I) APB northern blot analysis of queuosine on tRNA^Asp^, tRNA^Asn^, tRNA^His^ and tRNA^Tyr^ at 5.5 g/L (STD), 35.5 g/L NaCl and 55.5 g/L NaCl. Queuosinylation levels are compared against the Q-deficient Δ*tgt* mutant.

To investigate this further, we performed gene expression analysis of Q biosynthetic genes, which did not reveal any significant differences in the expression of *queA*, *tgt*, or *queG* at high salinity conditions **(Figure 3H)**. In contrast, by means of APB northern blot analysis, we observed a clear reduction in Q modification levels on the full-length tRNA^Asp^, tRNA^His^, and tRNA^Tyr^ isoacceptors, respectively. Moreover, tRNA^Asn^ failed to produce a clearly detectable band, most likely due to its low abundance in the tRNA pool **(Figure 3I)**. These observations might stem from a variety of salinity stress-induced factors, such as increased degradation of Q-modified tRNA (QtRNAs) **(Supplementary Figure S1)** or reduced expression and/or functionality of Q modifying enzymes. In summary, salinity stress seems to promote structural modifications that enhance tRNA stability, whereas decreased Q levels for some tRNA isoacceptors implies a potential regulatory function during physiological stress.

Given that MS and qPCR revealed distinct responses for the Q modification pathway, we decided to perform a transcriptomic analysis to identify additional processes that are affected by high-salinity (35.5 g/L and 55.5 g/L NaCl) at the early growth phase (OD_600_ = 0.8) as compared to standard (STD) growth conditions (5.5 g/L NaCl). Principal component analysis (PCA) of DESeq2 normalised expression data revealed a clear separation between stressed and STD growth conditions along principal component (PC)1, accounting for 85 % of the variance **(Supplementary Figure S2B)**. Biological replicates clustered closely in each group, confirming a coherent transcriptomic response. Next, gene ontology (GO) enrichment analysis showed a consistent upregulation of several amino acid biosynthesis processes, including leucine, arginine, isoleucine, histidine and threonine in both stress conditions **(Supplementary Figure S3A, B)**. Additionally, processes associated with cellular stress responses, such as proteolysis, protein unfolding, heat shock response, etc., were overrepresented. Together, our results suggest that high-salinity stress leads to perturbations in protein homeostasis. Therefore, our assumption is that Q enzyme activity is compromised under high salinity, and such impairment may lead to reduced Q modification on tRNAs. Furthermore, we also noted that sodium and calcium ion transmembrane transport processes were downregulated **(Supplementary Figure S3A, B)**. This is consistent with high-salinity stress as a reduced production of ion channel proteins would limit the influx of ions to the cell (Gregory & Boyd, 2021).

### Isoacceptor-specific changes in Q modification constitute an adaptive response that furthers cold-active growth

To explore the temperature-dependent effect on tRNA modifications and its impact on bacterial growth and translation, we cultured cells at four temperatures; cold-active (0 °C and 5 °C), standard (15 °C), and high (25 °C). As expected, the highest growth rates were observed at 15 °C and 25 °C, whereas cold-active growth at 0 °C and 5 °C occurred at a much slower pace **(Figure 4A)**. This overall metabolic reduction was evident also in translation, where the level of actively translating polysomes **(Figure 4B)** was substantially lower—but not abolished—during cold-active growth **(Figure 4B, C)**, suggesting that the cells undergo translational adaptation. Next, we characterized changes in tRNA modification levels relative to the standard growth temperature (15 °C), which uncovered a significant increase in methylated PTMs, such as m^5^U, m^7^G, and m^1^G **(Figure 4D)**, as well as a notable rise for Ψ and s^4^U modification levels **(Figure 4E)**. These PTMs are all located at the main body of the tRNA—except for m^1^G, which occupies position 37 adjacent to anticodon. These changes likely showcase an adaptive strategy to stabilize the tRNA structure and maintain translational activity across fluctuating temperature conditions. Interestingly, Q modification levels exhibited a pronounced 2-fold increase at 0 °C, whereas no significant change was detected at 5 °C or 25 °C **(Figure 4F)**. Furthermore, oQ levels were consistently upregulated at both cold (0 °C and 5 °C) and high (25 °C) temperatures **(Figure 4G)**. To validate these findings, we performed APB northern blots that showed an isoacceptor-specific response to cold-active growth, with tRNA^Asp^ displaying significantly higher Q modification levels than tRNA^His^ or tRNA^Tyr^. Conversely, a similar dichotomy was observed at 25 °C, although here the highest Q levels were detected for tRNA^Tyr^ and the lowest for tRNA^Asp^, respectively **(Figure 4H)**. Consistent with previous observations, tRNA^Asn^ did not yield a strong signal and no discernible differences were noted **(Figure 4H)**. Moreover, an RT-qPCR fold-change analysis of the queuosine biosynthesis genes *queG*, *tgt*, and *queA* revealed no major differences in transcript levels, indicating that the observed Q profiles arise through mechanisms distinct from transcriptional regulation **(Supplementary Figure S4)**. Overall, these results demonstrate that Q modification is not uniformly distributed across applicable tRNAs but is instead modulated as a function of temperature and isoacceptor identity.

**Figure 4.**
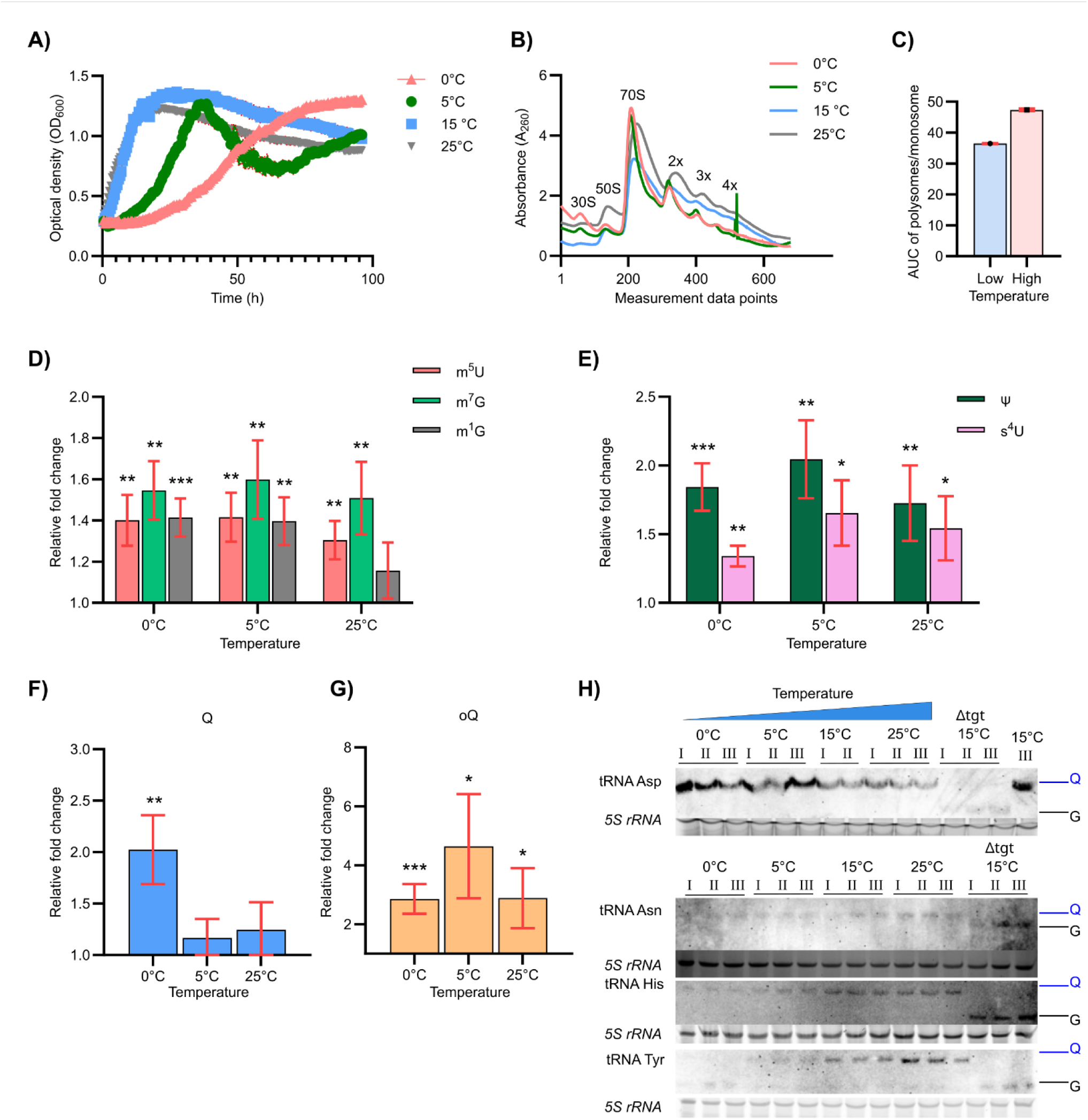
Temperature-induced changes to the *S. glacialimarina* tRNA modification landscape. A) Growth of *S. glacialimarina* at temperature conditions ranging from 0 °C to 25 °C. The shaded area denotes the standard deviation within the curve. B) Polysome profiles of *S. glacialimarina* following 10–40 % sucrose gradient centrifugation. The absorbance (A_260_) was monitored throughout the fractionation process and plotted as a chromatogram. The peaks corresponding to prokaryotic 30S, 50S, 70S rRNA along with monosome and polysomes are shown in the plot. C) Bar plot representation of the translation rate (determined as the ratio of the combined area under the curve (AUC) for all polysome peaks vs. the monosome peak) during cold-active (group Low, 0 °C and 5 °C) and high temperature (group High, 15 °C and 25 °C) growth. D-G) Bar plot of selected PTM modifications with significant trends observed at 0 °C, 5 °C, and 25 °C. PTM levels are presented as fold change relative to control (15 °C). Error bars denote standard deviation within the sample size n = 4 and *p*-values were calculated using one-sample Student’s *t*-test with significance represented as * *p* < 0.05, ** *p* < 0.005, *** *p* < 0.0005. H) APB northern blot analysis of Q on tRNA^Asp^, tRNA^Asn^, tRNA^His^, and tRNA^Tyr^ at 0 °C, 5 °C, 15 °C, and 25 °C. Queuosinylation levels are compared against the Q-deficient Δ*tgt* mutant.

To further assess transcriptional regulation at both temperature extremes for *S. glacialimarina*, we performed a differential gene expression analysis and compared DESeq2-normalised transcript abundances across the growth temperatures. PCA analysis showed that samples from 0 °C, 5 °C, and 15 °C formed distinct clusters, whereas the 25 °C samples grouped into two subclusters. These subclusters were positioned on the opposite side of PC1, indicating substantial variability within this condition **(Supplementary Figure S2A)**. Given that 25 °C is at the limit of sustainable growth for *S. glacialimarina* (Qasim *et al*., 2021), this variation likely reflects divergent adaptive responses. Despite this, we retained the 25 °C dataset for initial analysis but were cautious with our interpretation of trends arising from this condition. GO enrichment analysis revealed significant overrepresentation of upregulated fatty acid biosynthesis transcripts at 0 °C conditions, consistent with the established cold-adaptation strategy in psychrophilic bacteria maintaining membrane fluidity (Qasim *et al*., 2021) **(Supplementary Figure S5A)**. Additional upregulated processes at 0 °C included sodium ion transport, protein refolding, and multiple amino acid biosynthesis processes, such as leucine, arginine, histidine, and tryptophane. In contrast, transmembrane transport was the most downregulated category, along with processes related to the amino acid, potassium ion, and peptidoglycan transport. Similarly, upregulated processes at 5 °C included protein maturation, protein refolding, and chaperone co-factor dependent protein refolding, suggesting that sustained protein homeostasis is required at 5 °C to maintain metabolic activity. Conversely, genes involved in translation, ribosome biogenesis, and ribosome disassembly were downregulated **(Supplementary Figure S5B)**. However, these process categories included five or less gene transcripts at 25 °C, which limited our ability to infer clear functional trends **(Supplementary Figure S5C)**.

To extend our findings on isoacceptor-specific regulation of Q modification during temperature stress, we quantified the expression levels of individual isoacceptors in both the cold-active and high temperature groups. This analysis was designed to determine differences in Q abundance within the context of changes in the overall tRNA pool. PCA generated from DESeq2-normalised read counts grouped samples from each temperature group into distinct clusters **(Supplementary Figure S6A, B)**. Quality assessment using the tRNA-seq pipeline showed that > 90 % of all reads aligned uniquely to the tRNA reference sequences **(Supplementary Figure S7A)** and exhibited uniform coverage across the tRNAs **(Supplementary Figure S7B)**. Next, we quantified the relative abundance of Q-modified tRNA transcripts and assessed how their levels differed within the tRNA pool in the cold-active and high temperature groups. At cold-active temperatures, log_2_ fold change (FC) expression levels revealed a ≈2-fold decrease in tRNA^Tyr^ isoacceptor abundance **(Figure 5A-D, Supplementary Figure S8)**, which in part explains the lower Q modification levels observed for tRNA^Tyr^ **(Figure 4H)**. Interestingly, tRNA^Asp^ exhibited a ≈0.5-fold decrease in expression; yet Q modification levels increased sharply, suggesting a marked shift towards Q modified tRNA^Asp^ in the tRNA pool **(Figure 4H)**. In contrast, tRNA^Asn^ showed an ≈1-fold increase in expression **(Figure 5A-D)** with no corresponding change in Q modification detected by APB northern blot **(Figure 4H)**. Moreover, tRNA^His^ displayed no statistically significant difference in expression between the temperature groups, suggesting that the small decrease in Q modification observed at cold-active temperatures cannot be attributed to tRNA^His^ abundance **(Figure 5A-D**, **Figure 4H)**. Consequently, these results suggest that isoacceptor-specific Q modification changes cannot be solely driven by tRNA abundance. Therefore, increased Q modification on tRNA^Asp^ is likely to constitute an adaptive response by which *S. glacialimarina* drives the translation of genes important for its survival during cold-active growth. Similarly, the reduction in tRNA^Tyr^ abundance and its queuosinylation, together with a subtle decrease in Q on tRNA^His^, could be an additional layer of adaptation to cold stress. Furthermore, our analyses showed that tRNAs transcript expression patterns form one cluster at 0 °C and 5 °C (cold-active) and a second cluster at 15 °C and 25 °C **(Supplementary Figure S6A, B)**, supporting the hypothesis that the tRNA pool can reorganize under distinct physiological stresses to facilitate cellular stress adaptation (Torrent *et al*., 2018).

**Figure 5.**
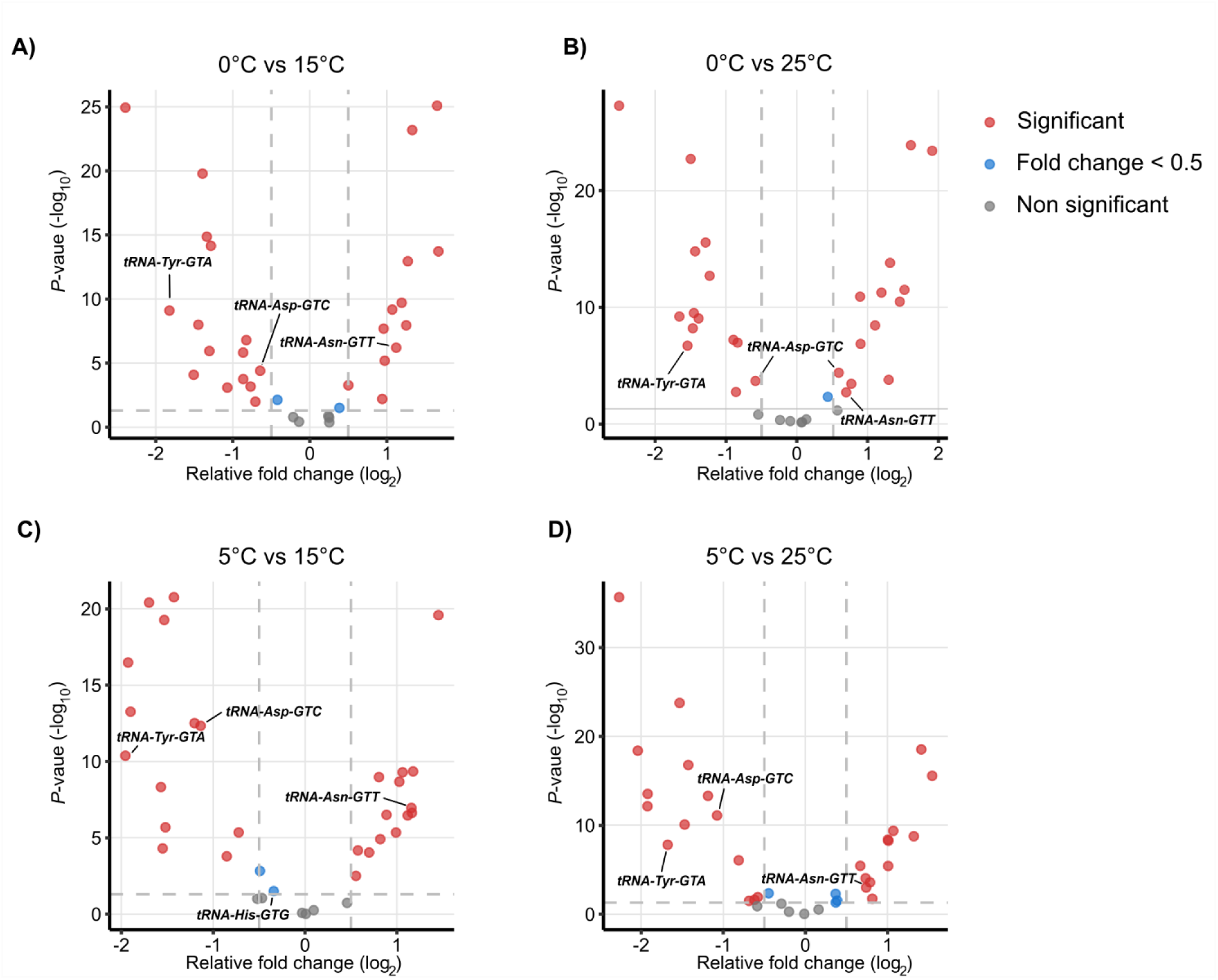
Comparison of temperature-driven expression level changes for Q-modifiable tRNA isoacceptors. Volcano plot of log2 fold change expression levels of tRNA molecules at A) 0 °C vs. 15 °C, B) 0 °C vs. 25 °C, C) 5 °C vs. 15 °C, and D) 5 °C vs 25 °C. The *y*-axis represents significance with −log_10_ adjusted *p*-value in each plot within a sample size of n = 4. Significantly up- or downregulated tRNA isoacceptors are denoted in red, < 0.5 log_2_FC are in blue, and non-significant changes are shown in grey.

### Queuosine-deficiency reduces the decoding ability of tRNA^His^ and triggers stress response pathways

To further characterize the effect of Q modification during cold-active growth, we performed comparative growth assays of the *S. glacialimarina* wild-type (WT) and Q modification deficient Δ*tgt* mutant at the aforementioned temperatures. This revealed a clear growth retardation for the Δ*tgt* mutant at cold-active temperatures, particularly at 0 °C **(Figure 6 A)**, whereas its growth rate was largely unaffected at 15 °C and 25 °C **(Supplementary Figure S9)**. To gain further insight into this phenotype, we proceeded with a comprehensive proteomic analysis that detected 2984 proteins (at 0 °C) and 2969 proteins (at 15 °C) out of 3725 predicted proteins. First, by calculating the log_2_FC between WT and Δ*tgt* at 0 °C and 15 °C **(Supplementary Data File 2)**, we verified that the *tgt* gene was successfully knocked out in the Δ*tgt* mutant **(Figure 6B)**, as was previously inferred by the loss of Q modification **(Figure 3I**, **Figure 4H)**. Moreover, we also observed a subtle polar effect (log₂FC ≈ 0.7_–_1 decrease) in the Δ*tgt* mutant for proteins expressed downstream of the *tgt* gene—i.e. the preprotein translocase subunit (YajC), protein translocase subunit (SecD), and protein translocase subunit (SecF) **(Supplementary Data File 2)**—which likely arises from the CRISPR gene deletion. Next, examining the Δ*tgt* dataset revealed a clear temperature-dependent upregulation of histidine biosynthesis proteins (HisF, HisH, HisC, HisD, HisA, HisG, and HisIE), where these proteins exhibited a log₂FC ≈ 1 at 15 °C, whereas this effect was almost log₂FC ≈ 2 at 0 °C **(Figure 6C)**. This observation, along with the reduction of Q modification on tRNA^His^ **(Figure 4H)**, suggests that the decoding efficiency of tRNA^His^ is reduced, thereby triggering ribosome stalling on the HisL leader peptide and subsequent activation of the histidine biosynthesis pathway. A similar response has been observed for *E. coli* and *V. cholerae*, where Δ*tgt*-mediated loss of Q modification induced histidine biosynthesis (de Crécy-Lagard *et al*., 2025).

**Figure 6.**
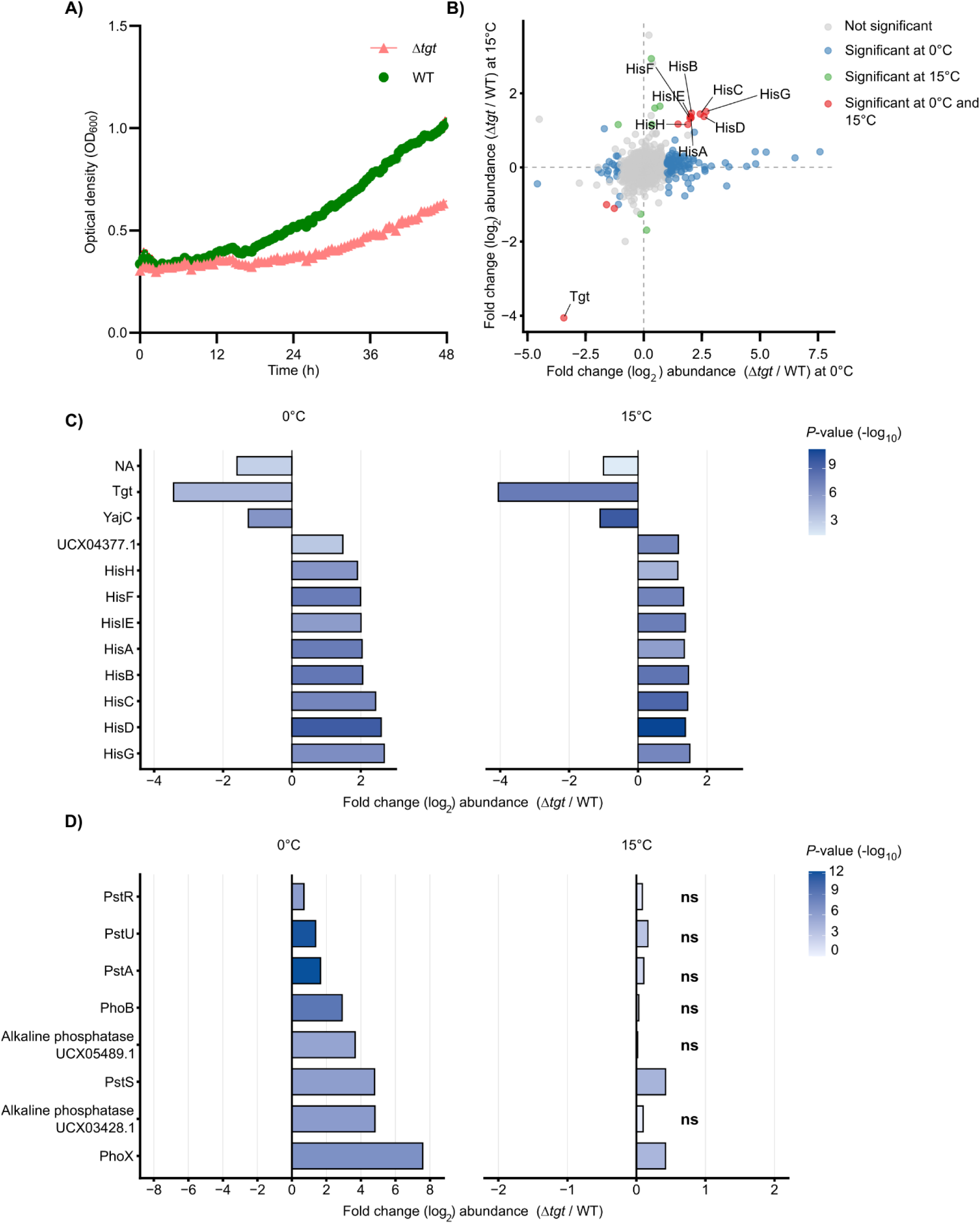
Loss of Q modification affects bacterial growth and translation. A) *S. glacialimarina* growth assays at 0 °C for WT strain and Δ*tgt* mutant (n = 5). B) Correlation plot of differentially expressed proteins in Δ*tgt*/WT at 0 °C and 15 °C, respectively (n = 5). Proteins with significant expression changes (Δ*tgt*/WT) at both 0 °C and 15 °C are denoted with red dots, while blue and green dots represent proteins with significant expression changes only at 0 °C or 15 °C, respectively. Proteins with non-significant changes are shown as grey dots. C) Horizontal bar plot of upregulated histidine biosynthesis proteins in Δ*tgt* at 0 °C and 15 °C (n = 5). D) Horizontal bar plot of upregulated phosphatase and phosphate transporter and regulatory proteins in Δ*tgt*/WT at 0 °C and 15 °C. For C) and D), protein symbols are on the *y*-axis and the light to dark blue gradient denotes significance (−log_10_ adjusted *p*-value, sample size n = 5). Full protein names are used if there are no established abbreviations, whereas protein ID is shown for unknown protein products.

In the absence of Q, guanosine pairs strongly with cytidine but weakly with uridine, creating a codon bias (Grosjean *et al*., 1978). Q modification mitigates this bias by stabilizing wobble pairing between C- and U-ending codons (Meier *et al*., 1985; Morris *et al*., 1999; Grosjean & Westhof, 2016). Based on this, we hypothesize that elevated Q levels on tRNA^Asp^ could influence the translation of genes important for adaptive responses during cold-active growth. Relative synonymous codon usage (RSCU), which compares codon frequencies assuming equal usage of synonymous codons, revealed a strong bias toward U-ending codons (NAU) **(Supplementary Figure S10)**. To assess whether this bias influence translation, we computed the standardized codon usage bias (SCUB) for all coding sequences, which normalises codon usage relative to the genome-wide pattern. Subsequently, we compared SCUB values for proteins significantly up- or downregulated at 0 °C in Δ*tgt* vs WT *S. glacialimarina* strains. This analysis revealed no significant difference **(Supplementary Figure S11)**, indicating that codon bias does not directly account for the observed changes in protein abundance at 0 °C. Moreover, functional analysis of the Δ*tgt* proteomics dataset at 0 °C showed a subtle but statistically significant increase for heat shock response pathways proteins (HSRs). For example, heat shock protein 20 (Hsp20) is increased by log₂FC ≈ 0.5 and aminoacyl-tRNA hydrolase is increased by log₂FC ≈ 1, as are several proteases, such as DegQ serine protease (log₂FC ≈ 0.35) and Clp protease (log₂FC ≈ 0.3). Indeed, this observation is consistent with activated protein quality control mechanisms **(Supplementary Data File 2)**, which suggests that loss of Q modification triggers stress response pathways to maintain protein homeostasis during cold-active growth.

### Queuosine-deficiency induces phosphate starvation in the *S. glacialimarina* Δ*tgt* mutant at 0 °C

While our data did not support codon-biased translation, it is well established that the decoding speed of synonymous codons determine the time window for accurate co-translational folding of nascent polypeptides (Nedialkova & Leidel, 2015). This intrinsic codon bias fine-tunes both the rate and fidelity of translation, thereby ensuring sufficient time for correct protein folding. For example, loss of m^1^G at tRNA position 37 disrupts the translation of membrane proteins with widespread consequences for cell envelope barrier integrity and efflux activity (Masuda *et al*., 2019), whereas aberrant modification of U_34_ results in loss of function of essential proteins and impairment of protein homeostasis (Nedialkova & Leidel, 2015). Hence, we further analysed the cold-active Δ*tgt* proteomics dataset for functional enrichment and observed a significant upregulation (log₂FC 1.3_–_4.7) for ion channel proteins, such as phosphate ABC transporter substrate-binding protein (PstS) and phosphate ABC transporter permease (PstA) and their regulatory proteins phosphate signalling complex protein (PhoU) and phosphate regulon transcriptional regulatory protein (PhoB) (**Figure 6D, Supplementary Data File 2)**. These proteins are a part of the Pho regulon (**Supplementary Figure S12A)** and facilitate the uptake of P_i_. Interestingly, the Δ*tgt* mutant at 0 °C also exhibited upregulation of alkaline phosphatases UCX03428.1 and UCX05489.1 (log₂FC ≈ 3.5) and a PhoX family phosphatase (log₂FC ≈ 7.5) **(Figure 6D, Supplementary Data File 2)**. At 15 °C, PstS and PhoX showed a modest increase in protein abundance (log₂FC ≈ 0.4), while PstR, PstU, PstA, PhoB, and the alkaline phosphatases exhibited no statistically significant differences. This suggests that the cells are experiencing phosphate starvation, an effect that is substantially amplified at 0 °C. While the underlying cause of this response is not fully understood, mistranslation of NAC/U codons or misfolding of proteins due to lack of Q is a plausible explanation.

## DISCUSSION

In this study, we present the first comprehensive tRNA epitranscriptome map of the cold-active marine bacterium *S. glacialimarina* TZS-4_T_. Using a multi-omics framework integrating genomic annotation predictions with experimental data from UPLC/MS and misincorporation sequencing (mim-tRNAseq), we rigorously characterized PTM levels in response to various physiological stressors. We observed a highly dynamic modification landscape that continuously fine-tunes to meet the translational demands of the cell. Notably, our temporal analysis revealed Q modification to be particularly responsive to bacterial growth, with the highest levels observed at the early exponential phase followed by a steady decrease until the 8 h timepoint. **(Figure 2B, C)**. Previous studies have shown that Q modification equilibrates the decoding of C- and U-ending codons, resulting in a stable translation rate (Meier *et al*., 1985; Morris *et al*., 1999; Tuorto *et al*., 2018). Therefore, a subtle increase in Q may facilitate protein translation during early exponential growth, thus generating a group of modification-tunable transcripts (MoTTs) (Gu *et al*., 2014). Yet in *S. glacialimarina*, the identification of MoTTs was hindered by the low GC content (41 %) of the genome. This creates a systematic bias towards A/U in the transcripts and increases the background level for NAU in coding sequences, effectively obscuring the signal after removing noise from the genome-wide bias. Nevertheless, it is widely regarded that Q on tRNAs is advantageous to ensure efficient and accurate decoding of NAU codons (Zaborske *et al*., 2014; Tuorto *et al*., 2018).

It is well established that tRNA PTMs can act as dynamic regulators of cellular physiology by rapidly adjusting the response to environmental cues, such as oxidative agents, temperature fluctuations, pH changes, antibiotics, toxins etc (reviewed in Teixeira *et al*., 2025). These PTMs are not merely a functional element of tRNA but also responsive regulators of cellular adaptation. Rather than constituting a uniform stress response, exposure to distinct stressors elicits a unique signature profile of PTMs. For instance, in *Saccharomyces cerevisiae* exposure to oxidative stress (H_2_O_2_) triggers upregulation of 5-methylcytosine (m^5^C), Cm, and N2,N2-dimethylguanosine (m^2,2^G), whereas exposure to alkylation agents like methyl methanesulfonate (MMS) or arsenite results in an entirely different modification landscape characterized by the modulation of wobble-uridine modifications, such as 5-methoxycarbonylmethyluridine (mcm^5^U) (Chan *et al*., 2010; Begley *et al*., 2007). In our study, we demonstrate that exposing *S. glacialimarina* to oxidative stress increases ROS and elicits significant changes on PTM levels, as seen for Q, thiolations, and methylations **(Figure 3A_–_C)**. This may be indicative of either diminished modification enzyme activity or rewiring of metabolic pathways to restore cellular growth. Conversely, high-salinity stress gave rise to a different response where modifications associated with the main body of the tRNA, such as s^2^C, s^4^U, Ψ, and acp^3^U, were upregulated **(Figure 3E, F)**. These adaptations may serve as a checkpoint for tRNA quality control by preventing degradation and promoting stability (Kimura & Waldor, 2019; Davis, 1995). Interestingly, we also observed a contradicting outcome for Q quantification by UPLC/MS and APB northern blot detection **(Figure 3G, H)**. We hypothesize that the elevated Q signal in the MS data may reflect salinity-induced tRNA degradation, whereby Q-containing fragments would remain detectable by MS, and/or perturbation of Q modification enzyme activity at high salinity **(Supplementary Figure S1).** Moreover, our observation of reduced Q across increasing salinity **(Figure 3I)** is more biologically meaningful for translation. A decline in Q levels could promote ribosome pausing, leading to disrupted proteostasis through greater protein misfolding, mistranslation, or frameshift errors (Tuorto *et al*., 2018). Consistent with this hypothesis, our transcriptomic analysis also revealed an induction of proteolysis-related and protein-unfolding genes, which is indicative of the bacterial unfolded protein response being triggered—and to which reduced Q levels may be a contributing factor **(Supplementary Figure S3)**.

In addition to the structural adaptation of tRNA and reduced levels of Q modification in oxidative and osmotic stress, we also investigated the impact of the growth temperature on PTM modulation, transcription and protein synthesis. As a cold-active bacterium, *S. glacialimarina* has diverse adaptive mechanisms to sustain growth at temperatures as low as 0 °C. Such cold-active growth strategies include promoting translation of polyunsaturated fatty acids to maintain membrane fluidity and produce cryoprotectants (such as glycerol and sucrose) and cold shock proteins (CSPs), thereby preventing freezing and the formation of undesired regulatory structures on mRNA (Jones *et al*., 1996; Phadtare & Severinov, 2010). Furthermore, cold-induced structural rigidity of tRNA is often counteracted by D modification, which alters base interactions by resisting base stacking and favouring the C2’-endo conformation, thereby increasing local flexibility (Dalluge *et al*., 1997). Accordingly, we show that several PTMs, particularly m^5^U, m^7^G, m^1^G, Ψ, and s^4^U—all located on the body of the tRNA—are divergently regulated at different temperatures in *S. glacialimarina* **(Figure 4D, E)**. Interestingly, at 0 °C we observed an isoacceptor-specific regulation for Q modification, where Q levels on tRNA^Asp^ are elevated whereas tRNA^His^ and tRNA^Tyr^ display reduced Q levels compared to standard conditions (15 °C) **(Figure 4H)**. This result suggests that tRNA^Asp^ is important for cold-active growth in *S. glacialimarina*, which was further confirmed by the growth retardation at cold-active temperatures for the Q-deficient Δ*tgt* mutant **(Figure 6A, Supplementary Figure S9A)**.

Next, we set out to explore how protein homeostasis shifts to enable cold-active growth. Using a comprehensive proteomics analysis, we uncovered that a key biological group of proteins associated with histidine biosynthesis were significantly upregulated in the *S. glacialimarina* Δ*tgt* mutant at 0 °C **(Figure 6C)**. Similar observations have been reported for *E. coli* and *Vibrio cholerae*, where the histidine biosynthesis proteins were upregulated in Δ*tgt* mutants (Pollo-Oliveira *et al*., 2022; de Crécy-Lagard *et al*., 2025). This histidine biosynthesis operon (**Supplementary Figure S12B)** is regulated by a leader peptide (HisL) characterized by its multi-His domain (Ames *et al*., 1983; de Crécy-Lagard *et al*., 2025). As previously described (Sherman *et al*., 1992), Q modification together with the tRNA discriminator position N73 serves as a key determinant for accurate aminoacylation. Thus, loss of Q is expected to decrease the pool of aminoacylated tRNA^His^. Simultaneously, Q-hypomodification on tRNA^His^ may also alter the decoding ability of the histidine codons. Together, these perturbation in aminoacylation and decoding can cause ribosome stalling on *hisL* leader sequence, triggering the expression of histidine biosynthesis genes (*hisGBDCBHAF*) **(Figure 7B)** to overcome the translation challenge imposed by impaired tRNA^His^.

**Figure 7.**
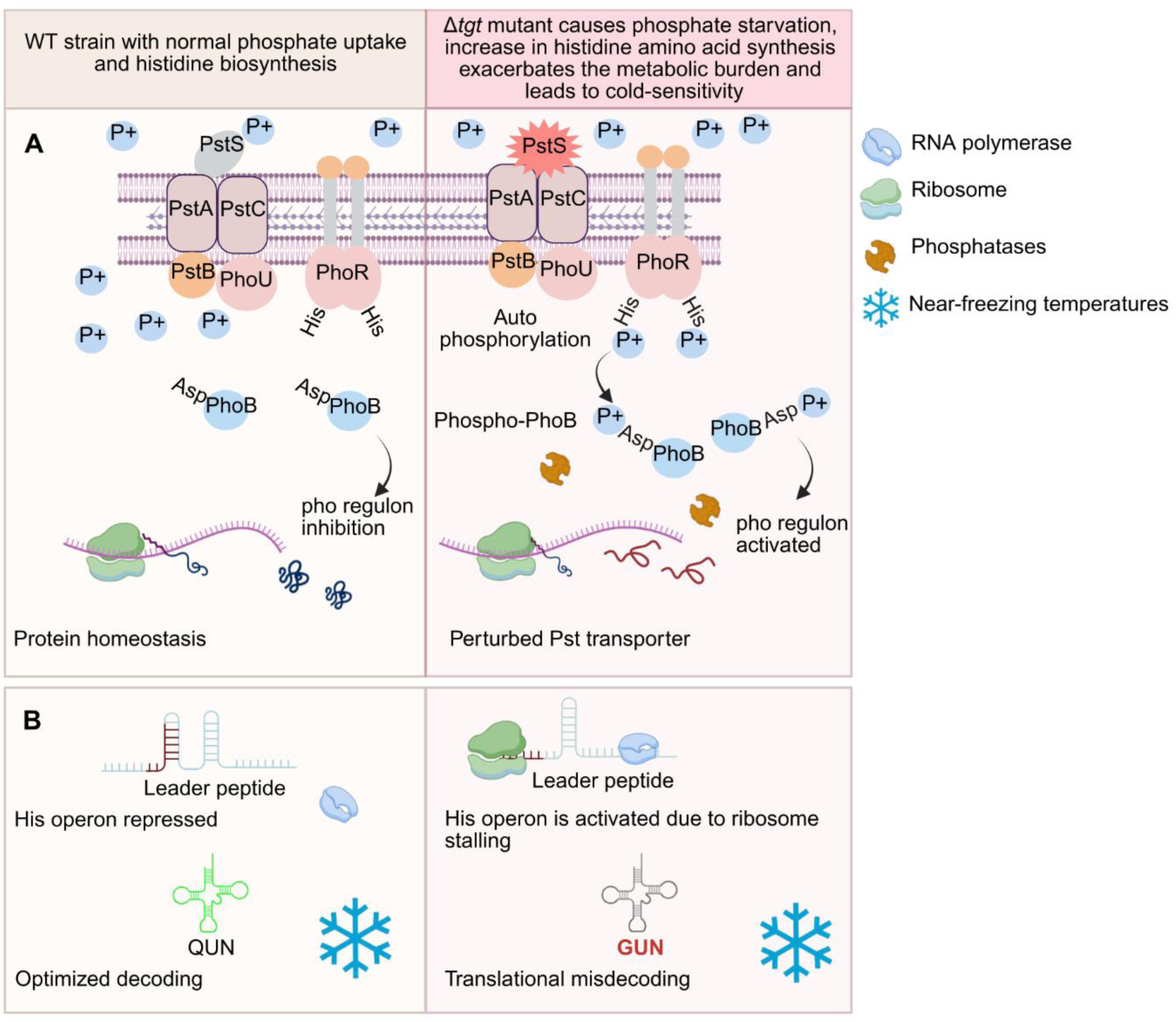
Proposed model illustrating how queuosine-deficiency contributes to cold-sensitivity in *S. glacialimarina*. A) Q modification-deficiency destabilizes protein homeostasis, manifesting as a phosphate-starvation phenotype. B) Hypomodified tRNA^His^ alters the decoding capacity and aminoacylation, triggering the histidine biosynthesis pathway. These concurrent stresses create a metabolic burden at near-freezing temperatures, which leads to cold-sensitivity and reduced growth. Created in BioRender.

Finally, we also examined the differential regulation of proteins in the context of transcript-wide codon usage bias **(Supplementary Figure S11)**. Due to the inherent GC bias of the *S. glacialimarina* genome, we were unable to identify codon-biased translation of MoTTs using SCUB analysis. However, RSCU analysis showed a clear overrepresentation of U-ending codons in the transcripts **(Supplementary Figure S10)**. Since Q maintains the translation rate in a context-specific manner by reducing the decoding bias between C- and U-ending codons (Grosjean *et al*., 1978; Tuorto *et al*., 2018), the loss of Q modification would decrease the translation efficiency of NAU codons and thereby systematically affect cold-active protein synthesis. Moreover, Q-deficiency also impacts the fidelity of translation, creating divergence in the protein pool that prompts the cell to compensate for potential loss of function by increasing cold-active protein synthesis—as observed for the phosphate ABC transporter protein PstS, which was 4-fold upregulated in the Δ*tgt* mutant **(Figure 6D)**.

Based on the results of this study, we propose a model for Q mediated cold-active growth in *S. glacialimarina* **(Figure 7)**. We hypothesize that perturbations in PstS translation may impair the activity of phosphate transporters and lead to phosphate starvation **(Figure 7A)**. Hence, a decrease in PstS activity reduces P_i_ uptake and creates a periplasmic phosphate insufficiency, which is recognized by the PhoR/PhoB two-component regulatory system (TCRS) in conjunction with the phosphate-specific transport (Pst) system and the PhoU regulatory protein. At standard growth conditions where P_i_ is abundant, the Pst system along with PhoU forms a repressive complex that inhibits PhoR autophosphorylation. In this inhibited state, PhoR acts as a phosphatase that—together with PhoU and PstB—actively dephosphorylates the response regulator PhoB, keeping the Pho regulon inactivated. Conversely, during P_i_ starvation, PhoR autophosphorylates histidine residues in its catalytic site **(Supplementary Figure S13)** and transitions into the active state. Once activated, PhoR transfers its phosphate to the PhoB aspartate residue located in its catalytic site **(Supplementary Figure S14)**. This in turn activates Phospho-PhoB that acts as a transcriptional regulator of the Pho regulon, controlling transcription of these genes to maintain phosphate levels (Wanner, 1996). Moreover, activation of the Pho regulon in the Δ*tgt* mutant also coincides with an increase in alkaline phosphatase and PhoX phosphatase expression **(Figure 6D)**. This most likely reflects a response for increased P_i_ uptake, although it may heighten the metabolic burden of the cell, thereby providing a plausible explanation for cold-sensitivity in *S. glacialimarina* **(Figure 6A)**. However, our proteomics analysis did reveal a subtle polar effect for YajC, SecD, and SecF, which are expressed downstream of the *tgt* gene. This most likely arises from the CRISPR knockout process and could potentially be a contributing factor to the observed growth retardation at cold-active temperatures. Nevertheless, studies on Δ*tgt* mutants in other bacteria have shown that the observed phenotype(s) overwhelmingly stem from the gene deletion itself and not from polar effect (Pollo-Oliveira et al., 2022; de Crécy-Lagard et al., 2025; Fruchard et al., 2025).

In conclusion, our integrated multi-omics study has uncovered a novel translation regulating function for Q modification during cold-active growth in *S. glacialimarina*. We demonstrate that isoacceptor-specific queuosinylation of tRNA^Asp^ is essential for growth at near-freezing temperatures, whereas Q-deficiency triggers histidine biosynthesis and phosphate starvation at 0 °C. This suggests that *S. glacialimarina* relies on Q-tRNAs to maintain stable decoding and efficient translation of NAC/NAU codons at cold conditions. These insights offer a framework for future studies on translational regulation and stress adaptation in cold-active microbes.

## MATERIALS AND METHODS

### Bacterial strains and growth conditions

*S. glacialimarina* strain TZS-4_T_ (GenBank accession: CP041216.1, DSM 115441) was cultivated as previous described (Qasim *et al*., 2021). Briefly, *S. glacialimarina* was grown on 25 % (w/v) rich marine broth (rMB) solid media plates (48 h, 15 °C) until the formation of visible colonies. For liquid starter cultures, 25 % rMB was inoculated with a single colony and grown for 48 h at 15 °C (except for 0 °C, where starter cultures were grown at 5 °C) with orbital shaking at 200 rpm. Fresh cultures were prepared into 25 % rMB and inoculated with the starter culture to obtain an optical density (OD_600_) of ≈ 0.2. The cultures were then grown until the desired optical density at different physiological conditions. The *S. glacialimarina* Δ*tgt* knockout mutant (Gregorova *et al*., 2026b) was grown as described above for TZS-4_T_.

### Growth assays

Growth of *S. glacialimarina* WT and Δ*tgt* mutant strains was assayed using a Bioscreen C (Oy Growth Curves Ab Ltd, Turku, Finland) microplate photometer in 25 % rMB media at different physiological conditions. Starter cultures were prepared similarly as described above. The cultures were grown from OD_600_ ≈ 0.2 in a honeycomb microplate with lids (Bioscreen, cat. no. 95025BIO) for 72 h. Growth curves were generated by plotting the OD_600_ values from each condition as a function of time.

### RNA isolation

Bacterial cells were harvested from 50 mL of liquid culture by centrifugation (3,200 × *g*, 10 min, 4 °C) and flash frozen in liquid nitrogen (stored at −70 °C). RNA was isolated from bacterial pellets using acidic-phenol extraction method (Chomczynski & Mackey, 1995; Qasim *et al*., 2021). The pellet was dissolved in 4 mL 0.9 % w/v NaCl followed by addition of 4 mL acidic phenol (Sigma-Aldrich, St. Louis, MO, USA, cat. no. P4682-400ML), 1 mL TRIzol and 800 µL of 1-bromo-3-chloropropane (Acros Organics, Geel, Belgium cat. no. 106860010). To facilitate cell lysis, glass beads were added to the lysate and vortexed vigorously for 10 min at room temperature (RT). The lysate was then centrifuged (10,000 × *g*, 10 min, RT) and RNA from the aqueous phase was precipitated with 99.6 % EtOH. The RNA pellets were air-dried and resuspended in RNase-free double distilled water (ddH_2_O). RNA concentration was determined using NanoDrop 2000 spectrophotometer (Thermo Scientific, Waltham, MA, USA) and resolved on a 1.2 % Tris-Borate EDTA (TBE) agarose gel, stained with Midori Green Advance (Nippon Genetics, cat. no. MG04; 4 µL per 100 mL of agarose solution) for assessment of RNA integrity. Gel images were captured on ChemiDoc Imaging System (Bio-Rad Laboratories, Hercules, CA, USA).

### Enrichment of transfer RNA using silica spin columns

Transfer RNA was enriched from total RNA using NucleoSpin RNA silica matrix containing columns (Macherey-Nagel, Düren, Germany cat. no. 740955.50S) as previously published (Gregorova *et al*., 2026a). Briefly, 200 µg of total RNA was resuspended in 100 µL of ddH_2_O and mixed with 250 µL of LBB (5 M guanidinium thiocyanate, 50 mM Tris pH 7.0). The suspension was mixed with 190 µL of binding buffer (100 % ethylene glycol diacetate, EGDA), vortexed, and loaded onto a NucleoSpin column. Following centrifugation (11,000 × *g*, 1 min, RT), the small RNA-containing flowthrough (FT) was collected. The FT (≈540 µL) was mixed with 675 µL of SBB (80 % ethylene glycol diacetate, 1 M guanidinium thiocyanate, 10 mM Tris pH 7.0) and thoroughly vortexed and incubated for 1 min. The mixture was then loaded onto a second NucleoSpin column and centrifuged (11,000 × *g*, 1 min, RT). The leftover FT was mixed with SBB as described above and reloaded onto the second column. The column was washed (11,000 × *g*, 1 min, RT) once with 500 µL of CW (1.6 M guanidinium thiocyanate, 66 % EtOH, 20 mM Tris pH 7.0) and twice with 500 µL of EWB (10 mM Tris-HCl pH 7.5, 20 mM NaCl, 80 % EtOH). The column was centrifuged (11,000 × *g*, 2 min, RT) and the purified tRNA was eluted with 50 µL of RNase-free ddH_2_O. The final concentration of the tRNA was estimated using NanoDrop 2000 spectrophotometer (Thermo Scientific) and stored at −70 °C.

### RNA sequencing and differential expression analysis

Total RNA was extracted from bacterial cells using the RNA isolation method described above. Ribosomal RNA (rRNA) depletion was performed using the RiboCop rRNA Depletion Kit for G- bacteria (Lexogen GmbH, Vienna, Austria; cat. no. 126.96) according to the manufacturer’s instructions. Sequencing libraries were prepared on the rRNA depleted samples using CORALL RNA-Seq V2 library prep kit with UDI 12 nt set A1 (Lexogen, cat. no. 171.96). The final libraries were amplified on a thermocycler (Avantor/VWR, Radnor, PA, USA. cat. no. 732-3428) using the cycle number estimated by qPCR. The sequencing libraries were pooled in equimolar proportions and sequenced on a NextSeq2000 (Illumina, Inc., San Diego, CA, USA) sequencer (100 cycles, single-end sequencing). Raw sequencing reads were demultiplexed using bcl2fastq (VBCF Next Generation Sequencing Facility) and quality control was performed using FastQC (v.0.12.1). Unique molecular identifiers (UMIs) were extracted using UMI-tools (v.1.1.5) extract command (Smith *et al*., 2017). Trimmomatic 3.39 with TrueSeq3-SE.fa file was used to remove the adapters with the parameter 2:30:10 (Bolger *et al*., 2014). Trimmed reads were aligned to its reference using BWA-mem (v.2.2) and sorted using samtools (v.1.19.2) (Li & Durbin, 2009; Li *et al*., 2009). The aligned bam files were deduplicated using UMI-tools command “umi_tools dedup”. Feature quantifications were conducted using HTSeq-count. Differential expression analysis was performed in R (v4.4.1) using DESeq2 (v1.44.0) with an adjusted *p*-value threshold of < 0.05 (Anders *et al*., 2015; Love *et al*., 2014).

### Gene ontology enrichment analysis

A custom organism annotation database for *S. glacialimarina* (TaxID: 2590884) was generated in R (v4.4.1) using the AnnotationForge (v.1.52.0) package. Genomic metadata were compiled from KEGG, with bacterial gene locus tags designated as primary identifiers. Annotation tables were formatted to meet *makeOrgPackage* specifications; GO terms were converted to the standard seven-digit format (GO:XXXXXXX), and all annotations were assigned the “Inferred from Electronic Annotation” (IEA) evidence code. The resulting SQLite-based package, *org.Sglacialimarina.eg.db*, was installed locally for subsequent functional analyses.

Gene Ontology (GO) enrichment analysis was performed using the topGO (v.2.56.0) package to identify over-represented biological processes. The gene universe was defined as all transcripts detected and differentially expressed genes (DEGs) were classified as up- or down-regulated based on an adjusted *p*-value < 0.05 and an absolute log_2_ fold change > 0.5. Enrichment analysis for the Biological Process (BP) ontology was performed using the Weight01 algorithm with Fisher’s exact test to account for hierarchical dependencies within the GO structure. Results were visualized using ggplot2 (v4.0.1) dot plots, where the “Rich Factor” (significant/annotated genes) denoted enrichment depth, and point size and colour represented gene counts and −log_10_ (*p*-value), respectively.

### Codon usage analysis

Standardized codon usage bias (SCUB) was calculated from coding sequences (CDS) using a custom R workflow. CDS files in FASTA format were imported with the Bioconductor package named Biostrings (v.3.19) and each sequence was parsed into codons according to the standard genetic code. Stop codons and amino acids represented by a single codon (methionine and tryptophan) were excluded because they lack synonymous alternatives. For each gene, codon counts were computed and normalised within each amino acid family to obtain gene-specific codon proportions. Genome-wide codon proportions were then calculated for each synonymous codon. Codon usage bias for each gene was defined as the difference between the gene-specific and genome-wide codon proportions. Bias values were standardized by dividing the codon-specific standard deviation across all genes, producing SCUB scores with approximately zero mean and unit variance. Codon usage patterns in differentially expressed genes were examined by filtering SCUB results using up- and down-regulated DEG lists, followed by visualization as violin plots using ggplot2 (v4.0.1). Relative codon usage analysis for all the coding sequences was calculated using a codon usage tool from The Sequence Manipulation Suite (Stothard, 2000).

### tRNA sequencing

Samples for tRNA sequencing library preparation were prepared as described (Pedor *et al.,* 2026) with minor modifications. Briefly, 200 ng of tRNA was purified on Monarch® Spin RNA Cleanup (10 µg) columns (New England Biolabs, Ipswich, MA, USA. cat. no. T2037-1) after diacylation. Next, each tRNA sample was linked with 3’ adapter containing unique indices (Behrens *et al*., 2022; Behrens & Nedialkova, 2022), indexed samples were pooled and reverse transcribed with in-house produced MRT-CBD (Pedor *et al*., 2026). The resulting cDNA was circularized, amplified using the Illumina TrueSeq small RNA barcoded adapters, and pooled into equimolar sequencing library. The sequencing was performed on a NovaSeqX 10B XP flowcell (Illumina) with paired-end sequencing and 150 cycles. The samples were demultiplexed using bcl2fastq (VBCF Next Generation Sequencing Facility). The sequencing data was analysed using mim-tRNAseq pipeline (Behrens *et al*., 2022; Behrens & Nedialkova, 2022) using the parameter; --no-cca --cluster-id 0.95 --threads 8 --min-cov 0.0005 --max-mismatches 0.1 --max-multi 4 --remap --remap-mismatches 0.075. The input file for the mim-tRNAseq pipeline was prepared using tRNAscan-SE 2.0.12 (Chan *et al*., 2021) and subsequently manually curated to conform to the GtRNAdb file format.

### Reverse transcription-quantitative PCR (RT-qPCR)

Target gene primers were designed using IDT PrimerQuest 2 tool and ordered from Metabion, Planegg, Germany. Total RNA (30 µg) was treated with 7 U of RQ1 RNase-free DNase (Thermo Scientific, cat. no. ENO521) and 3 µg of DNA-free RNA was reverse transcribed using 200 U of Maxima reverse transcriptase (Thermo Scientific cat. no. EP0742) primed with random hexamers (Qiagen, Hilden, Germany. cat. no. 79236; 0.2 µg/reaction). Both the DNase treatment and reverse transcription reaction were performed according to the manufacturer’s instruction. Primer specificity was validated using cDNA template amplification. The PCR products were analysed by electrophoresis on 1.2 % (w/v) tris-acetate-EDTA (TAE) agarose gels to confirm expected amplicon sizes. RT-qPCR reactions were performed using PerfeCTa SYBR green FastMix, Low ROX (Quantabio, cat. no. 95074-012). The reaction mix had 5 µL of 2× PerfeCTa mix, 0.4 µL of 2 µM forward and reverse primers with the final reaction volume set to 10 µL with ddH_2_O. Transcript amplification and quantification were performed on a Quantstudio 3 Real-time PCR system (Thermo Scientific) using the following PCR setup: 95 °C for 3 min followed by 50 cycles of 95 °C for 30 s, 60 °C for 30 s, and 72 °C for 30 s. Melting curve plots were generated to validate the formation of a single product. All the primers were tested for optimal amplification efficiency using the formula: primer efficiency (in %) = 100 x (10^−1/slope−1^) with 6-fold dilution of the template cDNA. Fold change analysis was performed using ΔΔC_T_ method (Livak & Schmittgen, 2001). Several housekeeping genes were previously tested, and gyrase A (*gyrA*) was kept as reference gene for the purpose of normalisation (Qasim *et al*., 2021). Data was analysed on QuantStudio design and analysis software (v1.5.2). All primer sequences corresponding to the genes and its amplification efficiencies are provided in **Supplementary Table T5**.

### Quantitative ultra-performance liquid chromatography-mass spectrometry (UPLC/MS) of tRNA modifications

Purified tRNA was digested and dephosphorylated into monoribonucleosides and separated on C18-UPLC as previously described (Gregorova *et al*., 2021). Chemical modifications on the ribonucleosides were validated by using commercially obtained modified ribonucleoside standards (Carbosynth Ltd., Compton, UK and Sigma). Peak identification and annotation of mass spectrometry data was implemented on MZmine4 (v.4.8.5) (Schmid *et al*., 2023) using a custom lookup list of modified ribonucleosides assembled from Modomics database and our in-house library of standard ribonucleosides (Gregorova *et al*., 2021). The signal intensities from each sample were normalised internally to adenosine (A), in addition to which the synthetic ribonucleoside 1,3-dimethylpseudouridine was used as a spike-in to verify consistent sample loading. The resulting quantification is presented as a relative change, defined as the ratio between normalised intensities of physiological stress condition and unstressed control samples.

### Assessment of Q levels in tRNA using APB northern blotting

The abundance of Q in tRNA^His^, tRNA^Tyr^, tRNA^Asp^, and tRNA^Asn^ was assessed using a non-radioactive biotinylated probe-based detection system on 3-acrylamidophenylboronic acid (APB) northern blots (Gregorova *et al*., 2026a). Briefly, 10 mL of APB bis-acrylamide solution (1× TAE, 7 M Urea, 5 % APB, 10 % bis-acrylamide [37.5:1]) was polymerized using 10 µL TEMED, 100 µL (NH_4_)_2_S_2_O_8_. Gels were hand-cast in the Mini-PROTEAN tetra vertical electrophoresis cell (Bio-Rad cat. no. 1658004) with 70 × 100 mm plates, 1 mm spacers, and a 16-well comb. Total RNA (4 µg) was deacetylated in 100 mM Tris-HCl (pH 9.0) and mixed with an equal volume of 2× RNA loading dye (90 % deionized formamide, 1× TAE, 0.05 % SDS, 0.01 % bromophenol blue) to a final volume of 20 µL. The RNA samples were denatured at 72 °C for 3 min and loaded onto a 10 % APB polyacrylamide gel. Electrophoresis was performed using 1× TAE as running buffer in a vertical chamber at 140 V and continued until the bromophenol blue dye front reached the bottom of the gel (approximately 2–3 h). Gels were stained with SYBR gold staining solution (10 µL SYBR in 50 mL 1× TAE) and visualized on Chemidoc (Bio-Rad). RNA was transferred onto an Amersham Hybond-N⁺ nylon membrane (Cytiva, cat. no. RPN203B) using 0.5× TBE as the transfer buffer in a semi-dry transfer cell (Bio-Rad, cat. no. 1703957). The membrane was crosslinked twice in a UV crosslinker CL-3000 (Analytik Jena GmbH, Jena, Germany) at 120 mJ/cm^2^. Pre-hybridization, probing, washing, and chemiluminescent detection steps were carried out as previously described (Gregorova *et al*., 2026a). Biotinylated probes for tRNA^His^, tRNA^Tyr^, tRNA^Asp^, and tRNA^Asn^ were synthesized by Metabion.

### Polysome profiling

*S. glacialimarina* cells were grown in 25 % rMB as described above. Cells were cultivated at 0 °C, 5 °C, 15 °C and 25 °C in 50 mL 25 % rMB until OD_600_ ≈0.8 was reached. The culture was flash-frozen and polysome profiling was carried out essentially as described (Mohammad & Buskirk, 2019). Briefly, bacterial cultures were sprayed into liquid nitrogen using a 50 mL serological pipette to form 7–12 mm (in diameter) ice pellets, which were mechanically pulverized in a prechilled grinding cylinder containing frozen lysis buffer (150 mM MgCl_2_) pellets. Each grinding cycle consisted of a 5 min pre-cooling phase followed by 5 grinding runs at 30 Hz for 1 min, and a 1 min cooling step after each grinding run. The pulverized cells were transferred to a pre-chilled 50 mL falcon tube. The samples were thawed on benchtop for 1–2 h (mix every 10 min) and placed on ice immediately afterwards. The lysates were pre-cleared by centrifugation (9,000 × *g*, 10 min, 4 °C) and a total of 22 mL of the pre-cleared supernatant was layered on top of 3 mL pre-chilled pelleting buffer (1.1 M sucrose, 20 mM Tris pH 8, 500 mM NH_4_Cl, 50 mM CaCl_2_, 4 % Triton X-100, 1 % NP-40, ddH_2_O) in two TH-641 tubes. Ribosomes were pelleted by centrifugation (205,835 × *g*, 3.5 h, 4 °C) using a T-865 fixed-angle rotor (Thermo Scientific), and the pellets were washed with 150 µL pre-chilled resuspension buffer (20 mM Tris pH 8, 15 mM MgCl_2_, 100 mM NH_4_Cl, 5 mM CaCl_2_, deionized H_2_O). Subsequently, each pellet was resuspended in 100 µL of resuspension buffer and combined. Next, 150 µL of ribosome resuspension (0.5 mg or 12.5 absorbance units ; [AU] 1 AU = A_260_ × 100/1000) was gently layered onto a 10–40 % sucrose gradient in an SW41 tube. The samples were centrifuged (201,000 × *g*, 2.5 h, 4 °C) using a TH-641 swinging bucket rotor (Thermo Scientific), after which they were fractionated in 300 µL fractions per tube using an automated gradient fractionator (BioComp Instruments, Tatamagouche, NS, Canada). The absorbance (A_260_) was measured for each fraction and plotted to generate a profile of ribosomal distribution across the gradient.

### Proteomics analysis with Liquid Chromatography–Electrospray Ionization–Tandem Mass Spectrometry (LC-ESI-MS/MS)

*S. glacialimarina* WT and Δ*tgt* mutant were grown in 25 % rMB to an OD_600_ ≈0.8 and samples were harvested by centrifugation at 3,200 × *g*. The pellets were dissolved in 90 µL TFA and incubated at 70 °C for 3 min. The samples were neutralized using 10 vol. of 2 M Tris-Base. Reduction and alkylation were performed by adding 99 µL of 10× reduction alkylation buffer (100 mM TCEP, 400 mM CAA) and incubated for 5 min at 95 °C. Samples were diluted with ddH_2_O water and digested overnight with sequencing-grade Trypsin/Lys-C (Promega, Madison, WI, USA). Peptides were desalted using a Sep-Pak tC18 96-well plate (Waters Corp., Milford, MA, USA), evaporated to dryness and stored at −20 °C until analysis. For LC-MS/MS analysis, peptides were reconstituted in 0.1 % (v/v) formic acid and analysed using a DIA analysis workflow on an EASY-nLC 1200 system coupled to an Orbitrap Exploris 480 (Thermo Scientific) mass spectrometer equipped with a nano-ESI source and FAIMS interface. Compensation voltages of −40 V and −60 V were applied. Peptides were loaded onto a trapping column and separated on a 15 cm C18 analytical column (75 µm × 15 cm, ReproSil-Pur 3 µm, 120 Å C18-AQ). Chromatography was performed using 60-min gradient (5–21 % solvent B over 28 min, 21–36 % over 22 min, 36–100 % over 5 min, followed by a 5 min wash). Solvent A consisted of 0.1 % formic acid in water, solvent B was 80:20 acetonitrile:water with 0.1 % formic acid. Wash runs were included between the samples. Data was acquired in DIA mode using Thermo Xcalibur 4.6 software. Each duty cycle included one full scan (400−1,000 m/z) followed by 30 DIA MS/MS scan using variable isolation windows across the same mass range.

Protein identification and label-free quantification were performed using Spectronaut (Biognosys, Schlieren, Switzerland; v.20.2). A DirectDIA workflow was applied for protein identification, and label-free quantification was carried out using MaxLFQ algorithm. The following parameters were used: Trypsin/P as the digestion enzyme, up to two missed cleavages, carbamidomethylation as a fixed modification, and acetylation (protein N-terminus) and methionine oxidation as variable modifications. Searches were performed against the *S. glacialimarina* protein database supplemented with universal protein contaminant database (Frankenfield *et al*., 2022). False discovery rate cutoff was 1 %. Quantification was based on MS2-level signals using the area under the curve within the defined integration boundaries. Signal intensities were normalised using retention-time-dependent regression (Callister *et al*., 2006). Differential abundance was assessed using unpaired Student’s *t*-test with the combined MS1 + MS2 statistical model (Huang *et al*., 2020). Multiple testing correction was performed using Storey’s approach to obtain *q*-values.

### Statistical analyses and data visualization

GraphPad Prism (v10.1.2) and R (v4.4.1; http://www.r-project.org) were used for all statistical analyses and visualizations. The statistical tests and biological replicate numbers are listed in each figure legend.

## DATA AVAILABILITY

All sequencing data generated in this study are publicly accessible through NCBI. Raw RNA-seq data have been deposited under BioProject PRJNA1426426 and tRNA-seq data have been deposited under BioProject PRJNA1425989.

## AUTHOR CONTRIBUTIONS

Conceptualization, M.S.Q. and L.P.S.; Data curation, M.S.Q.; Formal analysis, M.S.Q.; Funding acquisition, L.P.S.; Investigation, M.S.Q., N.S., and L.P.S.; Methodology, M.S.Q., J.K.P., A.W., J.M., O.K., N.S., and L.P.S.; Project administration, L.P.S.; Resources, L.P.S.; Supervision, L.P.S.; Validation, M.S.Q., J.K.P., A.W., J.M., O.K., and L.P.S.; Visualization, M.S.Q.; Writing – original draft, M.S.Q.; Writing – review and editing, M.S.Q., J.K.P, A.W., J.M., O.K., N.S., and L.P.S.

## DISCLOSURE AND COMPETING INTEREST STATEMENT

The authors declare that they have no conflict of interest.

## ACKNOWLEDGEMENTS

The authors thank Salla Kalaniemi and Sari Korhonen for their valuable technical assistance. The authors wish to acknowledge CSC - IT Center for Science, Finland for computational resources and the Next Generation Sequencing Facility at Vienna BioCenter Core Facilities (VBCF), member of the Vienna BioCenter (VBC), Austria, for NGS services. The facilities and expertise of the Instruct-HiLIFE Biocomplex unit at the University of Helsinki, a member of Instruct-ERIC Centre Finland, FINStruct, and Biocenter Finland are gratefully acknowledged. The authors are grateful to Miikka Olin, Department of Food and Nutrition, Faculty of Agriculture and Forestry, University of Helsinki for the use of their Synapt G2 Si mass spectrometer. Proteomics analysis was performed in the Turku Proteomics Facility, which is supported by Biocenter Finland. We also extend our gratitude to all members of the RNAcious laboratory for their insightful feedback and supportive discussions. This research was funded by the Research Council of Finland (grant #354906; to L.P.S.) and the Novo Nordisk Foundation (grant #NNF19OC0054454; to L.P.S.). M.S.Q. is a fellow of the Doctoral Programme in Microbiology and Biotechnology, University of Helsinki.

## REFERENCES

Allen EE, Facciotti D & Bartlett DH (1999) Monounsaturated but not polyunsaturated fatty acids are required for growth of the deep-sea bacterium *Photobacterium profundum* SS9 at high pressure and low temperature. Appl Environ Microbiol 65: 1710–1720

Ames BN, Tsang TH, Buck M & Christman MF (1983) The leader mRNA of the histidine attenuator region resembles tRNAHis: possible general regulatory implications. Proc Natl Acad Sci USA 80: 5240–5242

Anders S, Pyl PT & Huber W (2015) HTSeq—a Python framework to work with high-throughput sequencing data. Bioinformatics 31: 166–169

Begley U, Dyavaiah M, Patil A, Rooney JP, DiRenzo D, Young CM, Conklin DS, Zitomer RS & Begley TJ (2007) Trm9-Catalyzed tRNA Modifications Link Translation to the DNA Damage Response. Mol Cell 28: 860–870

Behrens A, Rodschinka G, Nedialkova DD. (2021) High-resolution quantitative profiling of tRNA abundance and modification status in eukaryotes by mim-tRNAseq. Mol Cell 81: 1802–1815.e7

Behrens A & Nedialkova DD (2022) Experimental and computational workflow for the analysis of tRNA pools from eukaryotic cells by mim-tRNAseq. STAR Protoc 3: 101579

Berger F, Morellet N, Menu F & Potier P (1996) Cold shock and cold acclimation proteins in the psychrotrophic bacterium *Arthrobacter globiformis* SI55. J Bacteriol 178: 2999–3007

Blaise M, Becker HD, Keith G, Cambillau C, Lapointe J, Giegé R & Kern D (2004) A minimalist glutamyl-tRNA synthetase dedicated to aminoacylation of the tRNAAsp QUC anticodon. Nucleic Acids Res 32: 2768–2775

Bolger AM, Lohse M & Usadel B (2014) Trimmomatic: a flexible trimmer for Illumina sequence data. Bioinformatics 30: 2114–2120

Callister SJ, Barry RC, Adkins JN, Johnson ET, Qian WJ, Webb-Robertson BJM, Smith RD & Lipton MS (2006) Normalization approaches for removing systematic biases associated with mass spectrometry and label-free proteomics. J Proteome Res 5: 277–286

Chan CTY, Dyavaiah M, Demott MS, Taghizadeh K & Dedon PC (2010) A Quantitative Systems Approach Reveals Dynamic Control of tRNA Modifications during Cellular Stress. PLoS Genet 6: 1001247

Chan PP, Lin BY, Mak AJ & Lowe TM (2021) tRNAscan-SE 2.0: improved detection and functional classification of transfer RNA genes. Nucleic Acids Res 49: 9077

Chomczynski P & Mackey K (1995) Substitution of Chloroform by Bromochloropropane in the Single-Step Method of RNA Isolation. Anal Biochem 225: 163–164

de Crécy-Lagard V, Baharoglu Z, Yuan Y, Boël G, Babor J, Bacusmo JM, Dedon PC, Ho P, Hummels KR & Kearns D (2025) Are bacterial processes dependent on global ribosome pausing affected by tRNA modification defects? J Mol Biol 437: 169107

de Crécy-Lagard V, Ross RL, Jaroch M, Marchand V, Eisenhart C, Brégeon D, Motorin Y, Limbach PA. (2020) Survey and Validation of tRNA Modifications and Their Corresponding Genes in *Bacillus subtilis* sp *subtilis* Strain 168. Biomolecules 10:977

Crick FHC (1966) Codon—anticodon pairing: The wobble hypothesis. J Mol Biol 19: 548–555

Dalluge JJ, Hamamoto T, Horikoshi K, Morita RY, Stetter KO & McCloskey JA (1997) Posttranscriptional modification of tRNA in psychrophilic bacteria. J Bacteriol 179: 1918–1923

D’Amico S, Collins T, Marx JC, Feller G & Gerday C (2006) Psychrophilic microorganisms: challenges for life. EMBO Rep 7: 385

Davis DR (1995) Stabilization of RNA stacking by pseudouridine. Nucleic Acids Res 23: 5020–5026

Endres L, Dedon PC & Begley TJ (2015) Codon-biased translation can be regulated by wobble-base tRNA modification systems during cellular stress responses. RNA Biol 12: 603–614

Frankenfield AM, Ni J, Ahmed M & Hao L (2022) Protein Contaminants Matter: Building Universal Protein Contaminant Libraries for DDA and DIA Proteomics. J Proteome Res 21: 2104–2113

Fruchard L, Babosan A, Carvalho A, Lang M, Li B, Duchateau M, Giai Gianetto Q, Matondo M, Bonhomme F, Hatin I, et al. (2025) Aminoglycoside tolerance in *Vibrio cholerae* engages translational reprogramming associated with queuosine tRNA modification. Elife 13

Gregorova P, Heinonen MMK, Laarne MM & Sarin LP (2026a) Protocol for rapid tRNA enrichment and chemiluminescent northern blot detection of tRNA and tRNA-derived fragments. STAR Protoc 7

Gregorova P, Heinonen M-MK, Sipari N & Sarin P (2026b) Queuosine promotes wecB-dependent phage resistance and biofilm formation in marine bacterium Shewanella glacialimarina. doi:10.64898/2026.03.05.709803 [PREPRINT]

Gregorova P, Sipari NH & Sarin LP (2021) Broad-range RNA modification analysis of complex biological samples using rapid C18-UPLC-MS. RNA Biol 18: 1382–1389

Gregory GJ & Boyd EF (2021) Stressed out: Bacterial response to high salinity using compatible solute biosynthesis and uptake systems, lessons from Vibrionaceae. Comput Struct Biotechnol J 19: 1014–1027

Grosjean H & Westhof E (2016) An integrated, structure- and energy-based view of the genetic code. Nucleic Acids Res 44: 8020–8040

Grosjean HJ, De Henau S & Crothers DM (1978) On the physical basis for ambiguity in genetic coding interactions. Proc Natl Acad Sci USA 75: 610–614

Gu C, Begley TJ & Dedon PC (2014) tRNA modifications regulate translation during cellular stress. FEBS Lett 588: 4287–4296

Huang T, Bruderer R, Muntel J, Xuan Y, Vitek O & Reiter L (2020) Combining Precursor and Fragment Information for Improved Detection of Differential Abundance in Data Independent Acquisition. Mol Cell Prot 19: 421–430

Johns GC & Somero GN (2004) Evolutionary Convergence in Adaptation of Proteins to Temperature: A 4-Lactate Dehydrogenases of *Pacific Damselfishes* (Chromis spp.). Mol Biol Evol 21: 314–320

Jones PG, Mitta M, Kim Y, Jiang W & Inouye M (1996) Cold shock induces a major ribosomal-associated protein that unwinds double-stranded RNA in *Escherichia coli*. Proc Natl Acad Sci USA 93: 76–80

Kimura S & Waldor MK (2019) The RNA degradosome promotes tRNA quality control through clearance of hypomodified tRNA. Proc Natl Acad Sci USA 116: 1394–1403

Lampi M, Gregorova P, Qasim MS, Ahlblad NC V & Sarin LP (2023) Bacteriophage Infection of the Marine Bacterium *Shewanella glacialimarina* Induces Dynamic Changes in tRNA Modifications. Microorganisms 11: 355

Li H & Durbin R (2009) Fast and accurate short read alignment with Burrows–Wheeler transform. Bioinformatics 25: 1754–1760

Li H, Handsaker B, Wysoker A, Fennell T, Ruan J, Homer N, Marth G, Abecasis G & Durbin R (2009) The Sequence Alignment/Map format and SAMtools. Bioinformatics 25: 2078–2079

Livak KJ & Schmittgen TD (2001) Analysis of relative gene expression data using real-time quantitative PCR and the 2−ΔΔ_CT_ method. Methods 25: 402–408

Love MI, Huber W, Anders S. (2014) Moderated estimation of fold change and dispersion for RNA-seq data with DESeq2. Genome Biol 15:550

De Maayer P, Anderson D, Cary C & Cowan DA (2014) Some like it cold: Understanding the survival strategies of psychrophiles. EMBO Rep 15: 508–517

Masuda I, Matsubara R, Christian T, Rojas ER, Yadavalli SS, Zhang L, Goulian M, Foster L, Huang KC & Hou YM (2019) tRNA Methylation Is a Global Determinant of Bacterial Multi-drug Resistance. Cell Syst 8: 302–314.e8

Meier F, Suter B, Grosjean H, Keith G & Kubli E (1985) Queuosine modification of the wobble base in tRNAHis influences ‘in vivo’ decoding properties. EMBO J 4: 823–827

Miles ZD, McCarty RM, Molnar G, Bandarian V. (2011) Discovery of epoxyqueuosine (oQ) reductase reveals parallels between halorespiration and tRNA modification. Proc Natl Acad Sci USA 108: 7368–72

Miyauchi K, Kimura S, Suzuki T. (2013) A cyclic form of N6-threonylcarbamoyladenosine as a widely distributed tRNA hypermodification. Nat Chem Biol 9:105–11

Mohammad F & Buskirk AR (2019) Protocol for Ribosome Profiling in Bacteria. Bio Protoc 9: e3468

Morris RC, Brown KG & Elliott MS (1999) The effect of queuosine on trna structure and function. J Biomol Struct Dyn 16: 757–774

Motorin Y & Helm M (2010) tRNA Stabilization by Modified Nucleotides. Biochemistry 49: 4934–4944

Nedialkova DD & Leidel SA (2015) Optimization of Codon Translation Rates via tRNA Modifications Maintains Proteome Integrity. Cell 161: 1606–1618

Neumann P, Lakomek K, Naumann PT, Erwin WM, Lauhon CT, Ficner R. (2014) Crystal structure of a 4-thiouridine synthetase-RNA complex reveals specificity of tRNA U8 modification. Nucleic Acids Res 42: 6673–85

Nichols DS, Nichols PD, McMeekin TA. (1993) Polyunsaturated fatty acids in Antarctic bacteria. Antarctic Science 5: 149–160

Ohira T & Suzuki T (2024) Transfer RNA modifications and cellular thermotolerance. Mol Cell 84: 94–106

Phadtare S & Severinov K (2010) RNA remodeling and gene regulation by cold shock proteins. RNA Biol 7: 788–795

Pollo-Oliveira L, Davis NK, Hossain I, Ho P, Yuan Y, Salguero García P, Pereira C, Byrne SR, Leng J, Sze M, et al. (2022) The absence of the queuosine tRNA modification leads to pleiotropic phenotypes revealing perturbations of metal and oxidative stress homeostasis in *Escherichia coli* K12. Metallomics 14: mfac065

Qasim MS, Lampi M, Heinonen M-MK, Garrido-Zabala B, Bamford DH, Käkelä R, Roine E & Sarin LP (2021) Cold-Active Shewanella glacialimarina TZS-4_T_ nov. Features a Temperature-Dependent Fatty Acid Profile and Putative Sialic Acid Metabolism. Front Microbiol 12: 737641

Russell NJ (1990) Cold adaptation of microorganisms. Philos Trans R Soc Lond B Biol Sci 326: 595–611

Salazar JC, Ambrogelly A, Crain PF, McCloskey JA & Söll D (2004) A truncated aminoacyl-tRNA synthetase modifies RNA. Proc Natl Acad Sci USA 101: 7536–7541

Schmid R, Heuckeroth S, Korf A, Smirnov A, Myers O, Dyrlund TS, Bushuiev R, Murray KJ, Hoffmann N, Lu M, Sarvepalli A, Zhang Z, Fleischauer M, Dührkop K, Wesner M, Hoogstra SJ, Rudt E, Mokshyna O, Brungs C, Ponomarov K, Mutabdžija L, Damiani T, Pudney CJ, Earll M, Helmer PO, Fallon TR, Schulze T, Rivas-Ubach A, Bilbao A, Richter H, Nothias LF, Wang M, Orešič M, Weng JK, Böcker S, Jeibmann A, Hayen H, Karst U, Dorrestein PC, Petras D, Du X, Pluskal T. (2023) Integrative analysis of multimodal mass spectrometry data in MZmine 3. Nat Biotechnol 41: 447–449

Sherman JM, Rogers MJ, Söll D. (1992) Competition of aminoacyl-tRNA synthetases for tRNA ensures the accuracy of aminoacylation. Nucleic Acids Res 20: 1547–52

Singh AK, Pindi PK, Dube S, Sundareswaran VR & Shivaji S (2009) Importance of *trmE* for growth of the psychrophile *Pseudomonas syringae* at low temperatures. Appl Environ Microbiol 75: 4419–4426

Smith T, Heger A & Sudbery I (2017) UMI-tools: Modelling sequencing errors in Unique Molecular Identifiers to improve quantification accuracy. Genome Res 27: gr.209601.116

Sordyl D, Boileau E, Bernat A, Maiti S, Mukherjee S, Moafinejad SN, Farsani MA, Shavina A, Cappannini A, Agostini G, Conticello SG, Stefaniak F, Dieterich C, Purta E, Bujnicki JM. (2026) MODOMICS: a database of RNA modifications and related information. 2025 update and 20th anniversary. Nucleic Acids Res 54: D219–D225

Stothard P (2000) The sequence manipulation suite: JavaScript programs for analyzing and formatting protein and DNA sequences. Biotechniques 28: 1102–1104

Suzuki T (2021) The expanding world of tRNA modifications and their disease relevance. Nature Reviews Mol Cell Biol 22: 375–392

Takakura M, Ishiguro K, Akichika S, Miyauchi K, Suzuki T. (2019) Biogenesis and functions of aminocarboxypropyluridine in tRNA. Nat Commun 10: 5542

Teixeira C, Vandenesch F & Moreau K (2025) tRNA modifications as regulators of bacterial virulence and stress responses. PLoS Pathog 21: e1013600

Torrent M, Chalancon G, De Groot NS, Wuster A & Madan Babu M (2018) Cells alter their tRNA abundance to selectively regulate protein synthesis during stress conditions. Sci Signal 11: eaat6409

Tuorto F, Legrand C, Cirzi C, Federico G, Liebers R, Müller M, Ehrenhofer-Murray AE, Dittmar G, Gröne H & Lyko F (2018) Queuosine-modified tRNAs confer nutritional control of protein translation. EMBO J 37: e99777

Urbonavičius J, Qian Q, Durand JMB, Hagervall TG & Björk GR (2001) Improvement of reading frame maintenance is a common function for several tRNA modifications. EMBO J 20: 4863–4873

Wanner BL (1996) Signal transduction in the control of phosphate-regulated genes of *Escherichia coli*. Kidney Int 49: 964–967

Watanabe K, Oshima T, Saneyoshi M & Nishimura S (1974) Replacement of ribothymidine by 5-methyl-2-thiouridine in sequence GTψC in tRNA of an extreme thermophile. FEBS Lett 43: 59–63

Watanabe K, Shinma M, Oshima T & Nishimura S (1976) Heat-induced stability of tRNA from an extreme thermophile, Thermus thermophilus. Biochem Biophys Res Commun 72: 1137–1144

Zaborske JM, DuMont VL, Wallace EW, Pan T, Aquadro CF, Drummond DA. (2014) A nutrient-driven tRNA modification alters translational fidelity and genome-wide protein coding across an animal genus. PLoS Biol 12: e1002015

